# Discovery of natural compounds as novel FMS-like tyrosine kinase-3 (FLT3) therapeutic inhibitors for the treatment of acute myeloid leukemia: an *in silico* approach

**DOI:** 10.1101/2024.08.27.609927

**Authors:** Uddalak Das, C Lavanya, Akshay Uttarkar, Jitendra Kumar, Vidya Niranjan

## Abstract

FLT3 mutations, observed in approximately 30-35% of Acute Myeloid Leukemia (AML) cases, drive leukemic proliferation and survival pathways, presenting a significant challenge in clinical management. To address this therapeutic need, we employed a comprehensive computational approach integrating pharmacophore screening, molecular docking, ADMET analysis, and molecular dynamics simulations to identify potent inhibitors targeting FLT3. Utilizing ligand-based pharmacophore models generated from experimentally proven FLT3 inhibitors from BindingDB, we screened over 400,000 natural compounds from the COCONUT database. Hits identified through pharmacophore screening underwent further evaluation via Lipinski and Golden triangle criteria to ensure drug-like properties. Molecular docking against the FLT3 receptor, combined with ADMET analyses, facilitated the prioritization of lead compounds. Subsequently, three promising candidates were subjected to molecular dynamics simulations to assess binding stability. Our findings reveal three top-performing compounds, demonstrating robust and stable binding affinity and favorable ADMET characteristics. These compounds hold promise as potential scaffolds or leads for developing novel FLT3 inhibitors in AML therapy.

## 1. INTRODUCTION

Acute myeloid leukemia (AML) represents a formidable hematologic malignancy characterized by aberrant myeloblast proliferation and impaired differentiation (Arber *et al*., 2016). With a prevalence exceeding 20,000 cases in the United States alone, AML is a significant healthcare burden, necessitating urgent exploration of novel therapeutic avenues (Cancer.Net, 2020; Majothi *et al*., 2020). Central to the pathogenesis of AML is the dysregulation of the FMS-like tyrosine kinase 3 (FLT3) gene, occurring in approximately 30% of newly diagnosed cases (Arber *et al*., 2016). AML patients frequently harbor mutations in FLT3, with two primary mutation types identified: internal tandem duplication (ITD) mutations, found in 25-35% of cases, and tyrosine kinase domain (TKD) mutations, observed in 5-10% of cases. These mutations confer constitutive activation to FLT3, leading to uncontrolled proliferation of leukemic cells and contributing to the aggressive nature of AML (Sakaguchi *et al*., 2019).

Despite advances in therapy, the prognosis for AML remains poor, with a five-year survival rate below 30% (Majothi *et al*., 2020). Targeted inhibition of FLT3 signaling has emerged as a promising therapeutic strategy, leading to the development of FLT3 inhibitors such as Midostaurin and Gilteritinib (Kiyoi *et al*., 2020). However, clinical efficacy is often hampered by the development of drug resistance, particularly in patients harboring FLT3-ITD/F691L mutations (Kawase *et al*., 2019). To address the challenge of drug resistance and improve therapeutic outcomes, there is a growing interest in the discovery of novel FLT3 inhibitors. Computational drug design approaches, including molecular docking and molecular dynamics simulations, offer a powerful means to expedite the identification and optimization of potential inhibitors (Mahmud *et al*., 2021). Understanding these interactions is essential for designing effective inhibitors with enhanced efficacy and reduced toxicity (Fernandes *et al*., 2020).

Despite these advancements, significant gaps remain in our understanding of FLT3-driven leukemogenesis and the development of effective targeted therapies for AML. While first- and second-generation FLT3 inhibitors have shown promise in clinical settings, challenges such as drug resistance and off-target effects persist, highlighting the need for innovative therapeutic approaches. Moreover, the potential of natural products in FLT3 inhibitor discovery, particularly in combination with computational methods and databases like COCONUT, remains underexplored.

In recent years, natural products have garnered attention as a valuable source of lead compounds for drug discovery. Their structural diversity and bioactivity make them attractive candidates for the development of novel therapeutics (Newman & Cragg, 2020). Integration of natural products with computational screening techniques presents an opportunity to identify potent FLT3 inhibitors from large compound libraries, such as the Collection of Open Natural Products (COCONUT) database (Sorokina *et al*., 2021).

In this study, we aim to bridge these gaps by comprehensively exploring the potential of natural products as a source of novel FLT3 inhibitors. By employing diverse ligand-based pharmacophore modeling, molecular docking, and molecular dynamics simulations, we seek to elucidate the molecular interactions between potential inhibitors and FLT3 protein residues. We endeavor to identify lead compounds with high binding affinity and selectivity for FLT3 through rigorous computational screening and analysis, offering promising candidates for further preclinical evaluation. By bridging the gap between computational drug design and natural product discovery, this study aims to contribute to the development of improved therapeutic strategies for AML, ultimately benefiting patients afflicted by this devastating disease.

## 1. MATERIALS AND METHODS

### 1.1. Protein preparation and receptor grid generation

In the Schrödinger Maestro protein preparation wizard, the protein was pre-processed with the PROPKA module for an optimization of H-bonds (Gokcan & Isayev, 2022), followed by minimization of structures towards convergence of heavy atoms at RMSD 0.3Å using OPLS4 force field and removal of water molecules more than 5Å away from ligands afterward (Mahdizadeh *et. al.,* 2021).

The receptor grid was generated keeping the hydrophobic region (predicted by SiteMap module) and also the region where Gilteritinib was attached to the complex, at the centroid of the grid. The coordinates of the receptor grid were X=-30.02, Y=-11.86, Z=-26.27, with ligand size upto 18 Å.

### 1.2. Dataset screening

COCONUT database (https://coconut.naturalproducts.net) was used as a database containing potential lead compounds. The following steps were employed to screen the compounds. At first, the compounds were screened by a generated pharmacophore model based on diverse ligands binding to FLT3. The screened hits were then prepared using the LigPrep module of Schrödinger. We employed the OPLS4 force field (Lu *et al.,* 2021), maintaining the ionization option as ‘Do not change.’ Standard procedures for desalination, tautomerization, and computational adjustments were conducted following software defaults. The prepared ligands were docked with the FLT3 (6JQR). Then, the pharmacokinetic parameters, specifically Lipinki’s Rule of 5 were considered to analyze and filter the compounds. Finally, the docked structures were subjected to molecular dynamics to analyze the interaction between hits and FLT3. The schematic illustrative diagram of the screening steps used in the study is depicted in Figure 3.

## 2. Pharmacophore modeling and screening

### 2.1.1. Pharmacophore hypothesis generation

To generate the pharmacophore model, the first 250 compounds related to the FLT3 gene were extracted from the binding database (https://www.bindingdb.org/bind/index.jsp) sorted by their respective IC50 values (Ganji *et. al.,* 2023). Four approved drugs that inhibit FLT3, namely Midostaurin (CID: 9829523), Quizartinib (CID: 24889392), and Gilteritinib (CID: 49803313) were used along with the relevant conformers for benchmark datasets This curated dataset was the foundation for subsequent pharmacophore model construction and screening endeavors. In the LigPrep module, the OPLS4 force field was used (Lu *et al.,* 2021), and the ionization option was set to “Do not change”. Furthermore, routine procedures involving desalination, tautomerization, and computational adjustments were implemented per software defaults. It helps to prepare high-quality 3D structures for drug-like molecules. Since Maestro format is the ideal and readable mode for Schrödinger software, this format was selected from the output directory.

In the Develop Pharmacophore Model module, the hypothesis match was set to 25%, as the dataset consisted of highly diverse active ligands. The number of features in the hypothesis was kept from 4 to 7 with the preferred number to 5. The ranking and ring of the hypothesis were set to the default “Phase Hypo Score” (Yu *et. al.,* 2021). The Generate Conformer and Minimize Output conformer options were activated, with the target number of conformers set to 50 (Cole *et. al.,* 2018).

### 2.1.2. Validation of the pharmacophore model

The parameters used for evaluating the efficiency of the developed pharmacophore model are enrichment factor (EF), receiver operating characteristic (ROC) curves (Triballeau *et al*., 2005), Boltzmann-enhanced discrimination of ROC (BEDROC) (Truchon & Bayly, 2007), and robust initial enhancement (RIE) (Truchon & Bayly, 2007).

A decoy set was created using the Generate DUDE Decoys program, which is found at http://dude.docking.org/generate (Mysinger *et. al.,* 2012). For converting the output into 3D structures, Open Babel v2.4.1 was used (O’Boyle *et. al.,* 2011). Ligand preparation was done following the default settings and the protocols of LigPrep. The OPLS4 force field (Lu *et al.,* 2021) has been employed in the minimization procedure.

### 2.1.3. Screening using the pharmacophore model

The best pharmacophore model “ADRR_2” based on BEDROC Score was used to screen the COCONUT database (Marondedze *et. al.,* 2020). 3 of 4 matches were selected and the “Prefer partial matches of more features” option was activated. The screening resulted in 83,429 compounds from the database of 4,07,270 compounds. After LigPrep, the screened compounds generated 3,46,926 structures.

#### 2.2. Molecular Docking Studies

Docking was limited to ligands with 100 rotatable bonds and fewer than 500 atoms. Van der Waals radii scaling factor was set to 0.80, with a partial charge cutoff of 0.15. Sample nitrogen inversions and sample ring conformations were activated, and the ligand sampling was set to flexible. All predicted functional groups had bias sampling of torsions enabled. The module was configured to promote intramolecular hydrogen bonds and improve conjugated pi groups’ planarity. Docking was done using the Glide module of Schrödinger.

Docking was done through a series of hierarchical filters i.e. HTVS mode (high-throughput virtual screening) for efficiently screening million compound libraries, to the SP mode (standard precision) for reliably docking tens to hundreds of thousands of ligands with high accuracy, to the XP mode (extra precision) where further elimination of false positives is accomplished by more extensive sampling and advanced scoring, resulting in even higher enrichment. Each step proceeded with the top 10% from the previous one. The HTVS filtered the library to around 1,97,852, SP filtered to around 23, 800 and, XP filtered to around 2,385 structures respectively

#### 2.3. *In silico* ADME/T and toxicity analysis

The docked ligands were screened based on “Golden Triangle” using the QikProp module of Schrodinger. Only compounds showing zero golden triangle violations were considered for molecular dynamics simulation studies. QikProp, SwissADME (Daina *et al.,* 2017), ProTox-II (Banerjee *et. al., 2018)*, and pkCSM (Pires *et. al.,* 2015) server were used in the analysis of pharmacokinetic properties to assess the detailed ADMET properties of those two compounds.

#### 2.4. Molecular dynamics simulation studies

Desmond package was used to carry out the molecular dynamics simulations for the FLT3- CNP0099279, FLT3-CNP0347183, and FLT3-CNP0298793 complexes. Each system was placed individually in an orthorhombic water box of 8Å using the SPC water model (Alturki *et. al.,* 2022). The ligand-protein complexes were modeled by the OPLS4 force field (Lu *et al*., 2021). Counter ions (Na+) were introduced in the ligand-protein complex structures to neutralize the total charge of the systems undergoing MD simulation. Furthermore, the energies of the systems were minimized to a minimum level using 2000 steps before initiating the MD simulation along an NPT lattice trajectory, for 100ns each and 310K temperature. (Al-Jumaili *et al.,* 2021).

The RMSD primarily suggests the stability of the ligand interaction, while RMSF describes the fluctuation and flexibility of the residues within the protein, particularly within the active site that is crucial for drug discovery. RMSD is calculated by the discovery 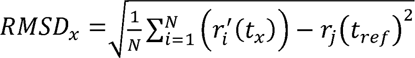. RMSF values also help in defining protein structures as they provide information about local conformational changes within the protein chains. (Bharadwaj, Bhargava, & Ball, 2021). The RMSF values are in units of Å and is calculated by the following equation 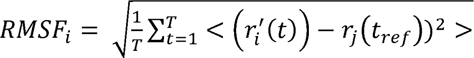

#### 2.5. Principal Component Analysis (PCA) and Dynamical Cross-Correlation Matrix (DCCM) Analysis

The trajectory file generated after the MD simulation was first extracted as a .cms file. This .cms file was then converted to .dcd format with the help of VMD 1.9.3 software (Humphrey *et al*., 1996). The trajectory .dcd file was subsequently uploaded to the MDM-TASK web server to perform PCA and DCCM analysis (Amamuddy *et al*., 2021). PCA was conducted to identify the major conformational changes and essential dynamics of the protein-ligand complex, while DCCM analysis was used to examine the correlated motions of residues over the simulation period.

#### 2.6. MM-GBSA analysis

The energy parameters generated by the MM-GBSA (Molecular Mechanics-Generalized Born Surface Area) simulation were predicted using the prime module. This was performed to predict the stabilization energy levels of potential interactions between the three selected ligands and the receptor. VGSB solvation model was used with OPLS4 force field (Hong *et. al.* 2024).

Using MD simulations that lasted 100ns, MM-GBSA (Molecular Mechanics-Generalized Born Surface Area) calculations to determine the binding free energy of protein-ligand complexes was done. This method of calculating binding energies proved more reliable than using glide scores obtained from molecular docking. The key energy components considered in the MM-GBSA calculations included hydrogen bond interaction energy, electrostatic solvation free energy, Coulomb (electrostatic) interaction energy, lipophilic interaction energy, and van der Waals interaction energy. These factors were all taken into account to estimate the relative binding affinity of the complexes.

## 3. RESULTS

### 3.1. The FLT3 Protein

The X-ray crystallographic structure of the FLT3 target protein [PDB ID: 6JQR], found at https://www.rcsb.org, in complex with Gilteritinib and determined to a resolution of 2.20Å by Kawase *et. al.* (2019), was used in the study.

Human FLT3 has a length of 993 amino acids and weighs 112.903 Da, having three components – the extracellular domain with 516 residues, the transmembrane domain with 19 amino acids, and the cytoplasmic domain of 429 amino acids (Ouassaf *et. al.,* 2023). To determine the structure, the PDBsum web server was utilized; this analysis identified fifteen α-helices, six β-hairpins, four β-bulges, as well as 12 helix-helix interactions. Furthermore, the Ramachandran plot was also used to validate FLT3, and it indicated that 93.5% of the residues were in preferred regions, 6.5% were in additional residue regions, none in generous regions, and 0.9% in disallowed regions with a total G-Factor of 0.00. (Ramachandran *et. al.,* 1963) (see Figure 2)

**Figure 1.**
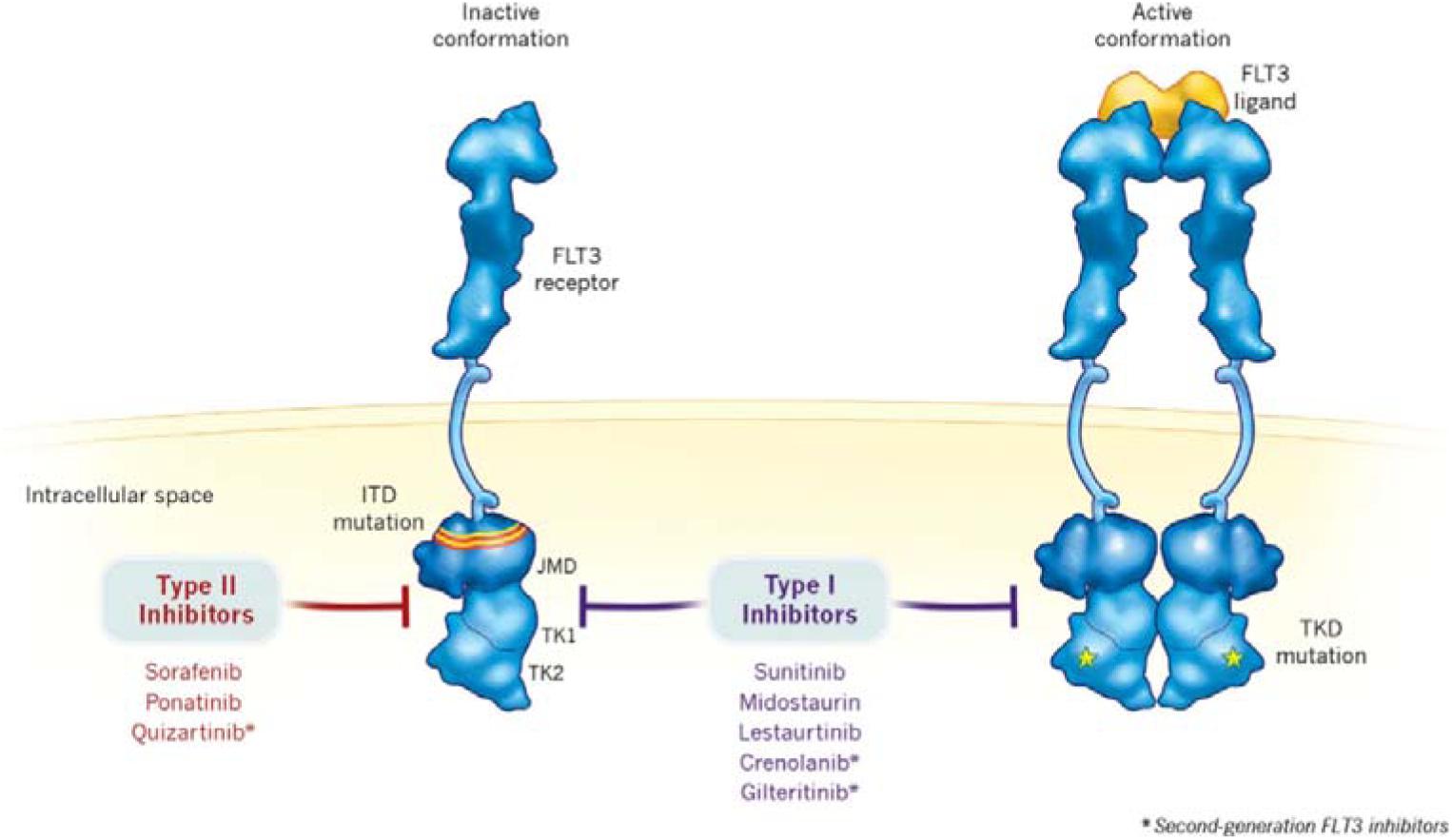
Type I FLT3 inhibitors attach to the active form of the FLT3 receptor, near the activation loop or ATP-binding pocket, and work against both ITD and TKD mutations. Type II FLT3 inhibitors bind to the inactive form of the FLT3 receptor, near the ATP-binding domain, specifically blocking ITD mutations but not TKD mutations. (Daver et. al., 2015) (Image source: Daver et. al., 2019)

**Figure 2.**
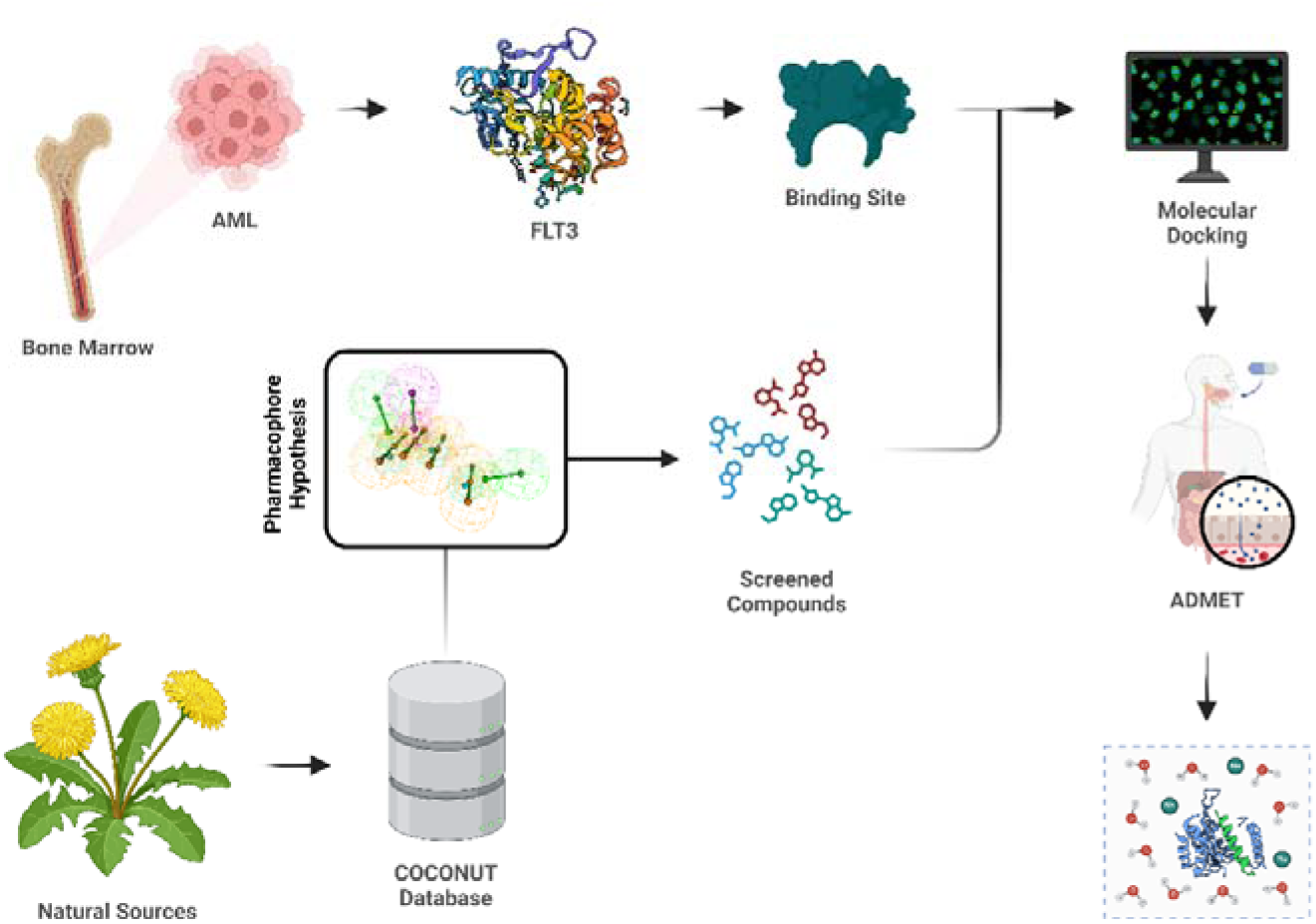
The schematic diagram of the complete dataset screening steps used in this study

**Figure 3.**
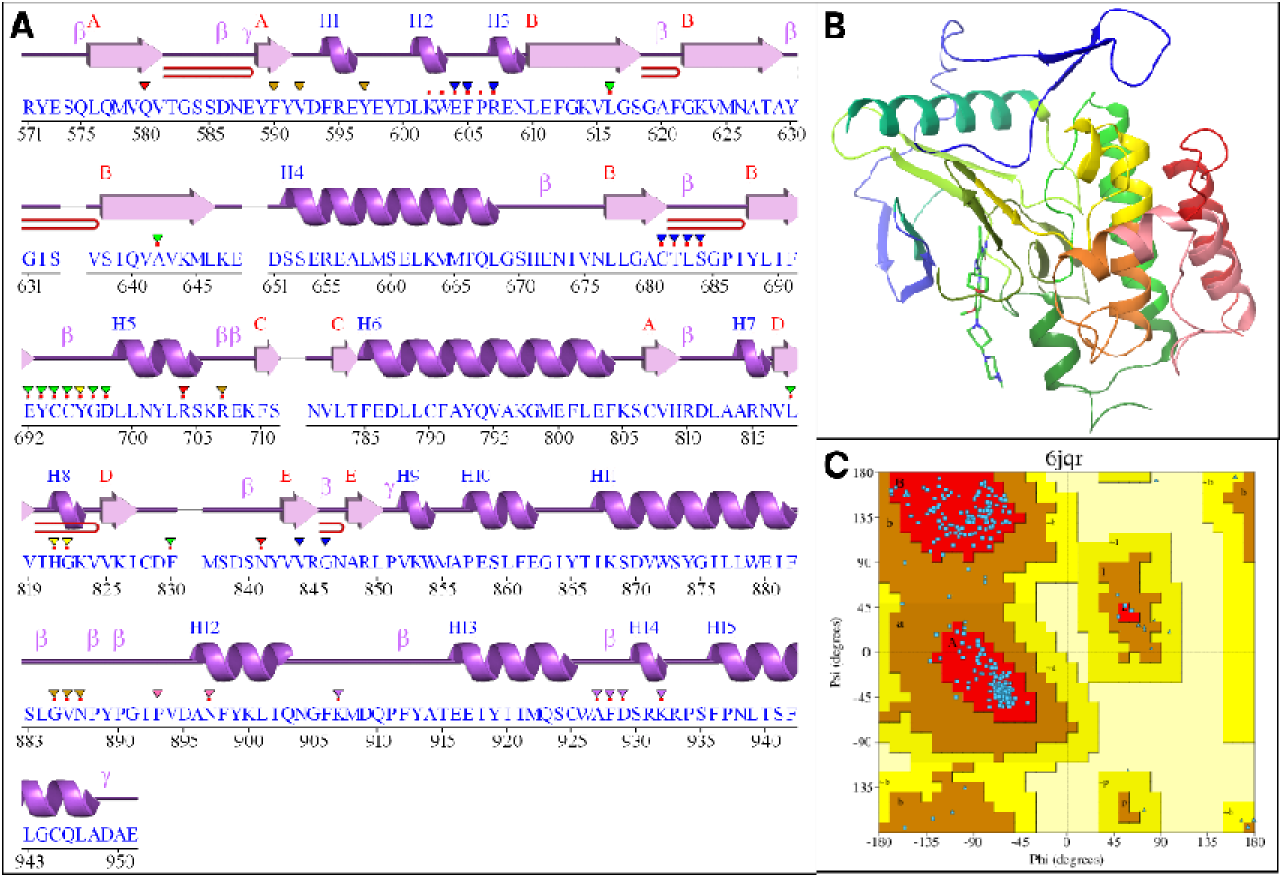
A: The secondary structure of the amino acid sequence of protein 6JQR (generated by PDBsum); B: The co- crystalized 3D structure of the FLT3 protein with its inhibitor Gilteritinib; C: Ramachandran plot of the protein. The color red indicates Iow-energy regions, yellow - allowed regions, pale yellow - generously allowed regions, and white - disallowed regions.

### 3.2. Pharmacophore modelling/screening

The generated pharmacophore models were ranked automatically based on the BEDROC Score. The hypothesis ADRR_2 revealed the highest PhaseHypo Score of 1.279276 comprising one hydrogen bond acceptor (A), one hydrogen bond donor (D), and two aromatic rings (R) features.

**Table 1.** Statistical values of different parameters of the top 10 generated pharmacophore hypothesis.

| Hypothesis ID | PhaseHypo<br>Score | Survival<br>Score | Site<br>Score | Vector<br>Score | Volume<br>Score | BEDROC<br>Score |
| --- | --- | --- | --- | --- | --- | --- |
| ADRR_2 | 1.279276 | 5.764613 | 0.722581 | 0.878267 | 0.630862 | 0.933367 |
| ADRR_1 | 1.278276 | 5.779649 | 0.730133 | 0.901594 | 0.619180 | 0.931503 |
| DRRR_1 | 1.268708 | 6.001781 | 0.727387 | 0.950273 | 0.662953 | 0.908568 |
| DRRR_2 | 1.266904 | 5.963407 | 0.717917 | 0.933409 | 0.650838 | 0.909149 |
| DHRR_2 | 1.253880 | 5.783011 | 0.549320 | 0.863629 | 0.534004 | 0.906941 |
| DHRR_1 | 1.247770 | 5.819535 | 0.569104 | 0.868776 | 0.564972 | 0.898633 |
| DRRR_4 | 1.244470 | 5.849485 | 0.683864 | 0.884103 | 0.621290 | 0.893550 |
| DRRR_3 | 1.241948 | 5.865835 | 0.693047 | 0.876708 | 0.634128 | 0.890004 |
| ADRR_3 | 1.237998 | 5.763253 | 0.602948 | 0.890596 | 0.605610 | 0.892228 |
| DDRRR_3 | 1.232596 | 6.256602 | 0.684735 | 0.882093 | 0.721721 | 0.857227 |

### 3.3. Pharmacophore Models Validation

The pharmacophore model developed was rigorously validated to assess its ability to accurately predict the activity of novel compounds identified through database screening or synthesized de novo. Validation is an essential step before utilizing a pharmacophore model for virtual screening (Kaserer *et al*., 2015).

The ADRR_2 model was applied to a test set database comprising 200 inactive molecules generated via the DUDE Decoys tool. It screened the decoy set to only 16 molecules indicating an efficiency of 92%. Further, validation of the ADRR_2 hypothesis revealed that EF in the top 1% of the decoy dataset is 3.43%, demonstrating that pharmacophore model is 6.43-fold efficient in detecting true positives/actives from the entire dataset. ROC score, RIE, and AUAC values were calculated as 0.63, 5.19, and 0.73, respectively. Thus, ADRR_2 was statistically significant in picking the actives from the decoy dataset. Statistical significance of model was also validated by calculating BEDROC. Contrary to EF, BEDROC seeks to measure the early enrichment of the actives. BEDROC values were calculated at different tuning parameter values (α[=[8.0, α[=[20.0, and α[=[160.9) and found to be 0.720, 0.844, and 1.000, respectively.

### 3.4. Molecular Docking Studies

The docking score range for the top 500 hits compounds was found between −19.195 to − 12.678 kcal/mol. The top 3 compounds based on docking score, with zero golden triangle violation, i.e., CNP0099279 (−16.041 kcal/mol), CNP0298793 (−14.869 kcal/mol), and CNP0347183 (−14.62 kcal/mol), were selected for further studies.

The ligand CNP0099279 showed diverse interactions with multiple ligands in the binding pocket of FLT3. It interacted with Asp829 with its H_2_N via H-bond, Glu661 with H_2_N and N^+^H_2_ vis H-bond and salt bridge respectively. It also interacted with Leu689 via H-bond with its OH and O, Ile687 with its HO via H-bond, Met625 with O via H-bond, and finally Lys623 via H-bonds with two OH groups. The ligand CNP0298793 showed interactions with 2 residues namely Ala627 and Lys623 via H-bonds with OH groups. CNP0347183 showed varying interactions with multiple residues viz. Met625 (H-bond with HN, NH, and O), Phe691 (H-bond with O), Leu689 (H bond with O and NH), and Glu661 (Salt Bridge with H_3_N^+^) (see Figure 5).

**Figure 4.**
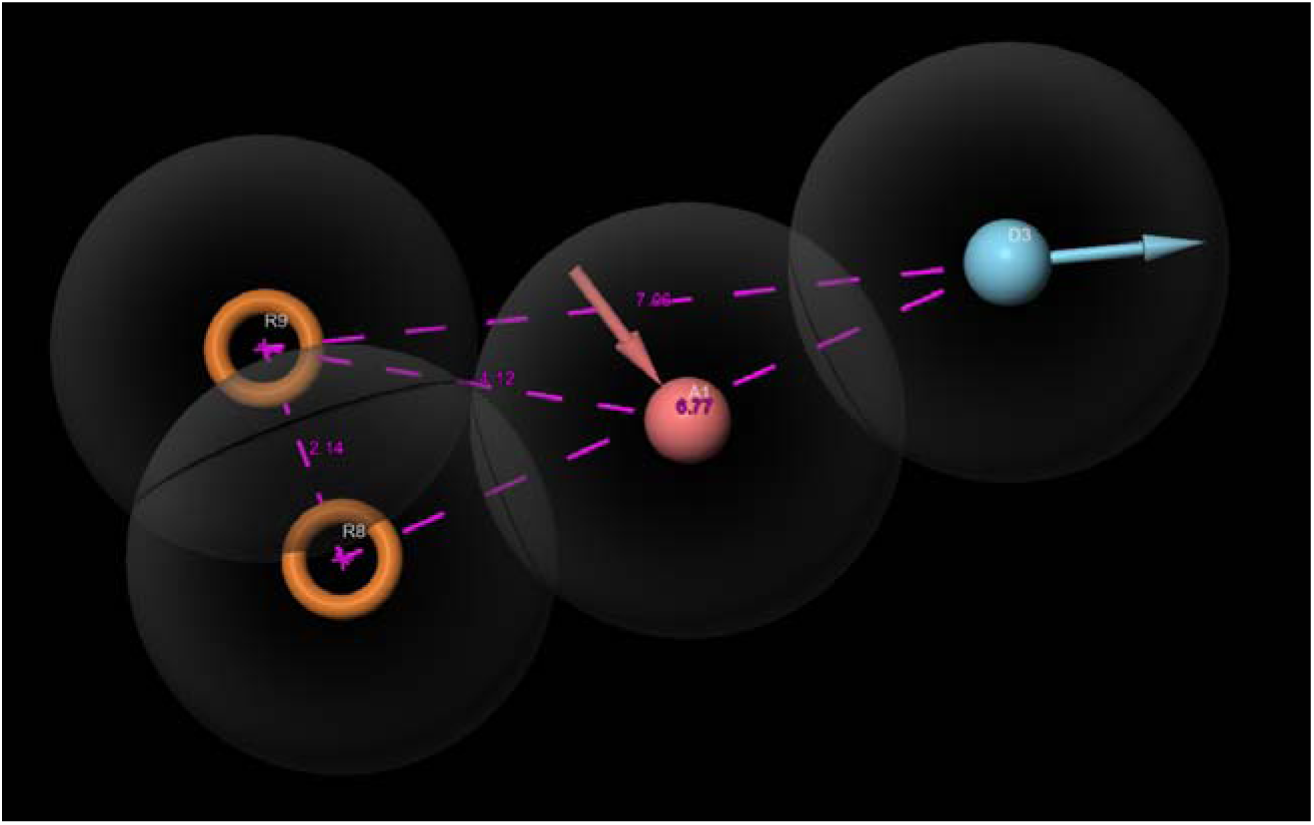
The distance between pharmacophore features of the best pharmacophore model, ADRR_2

**Figure 5.**
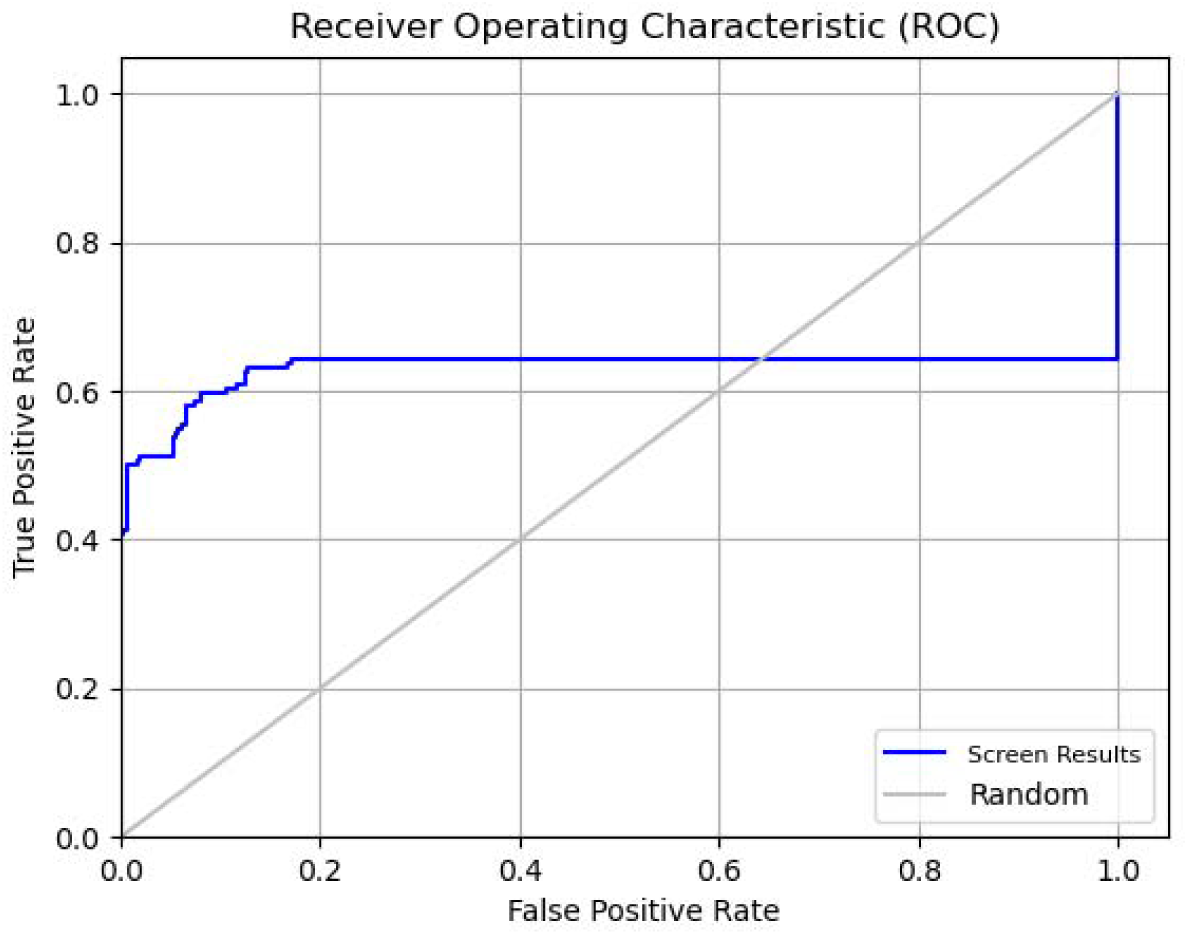
ROC curve represents the pharmacophore validation of the model ADRR_2

**Table 2.**
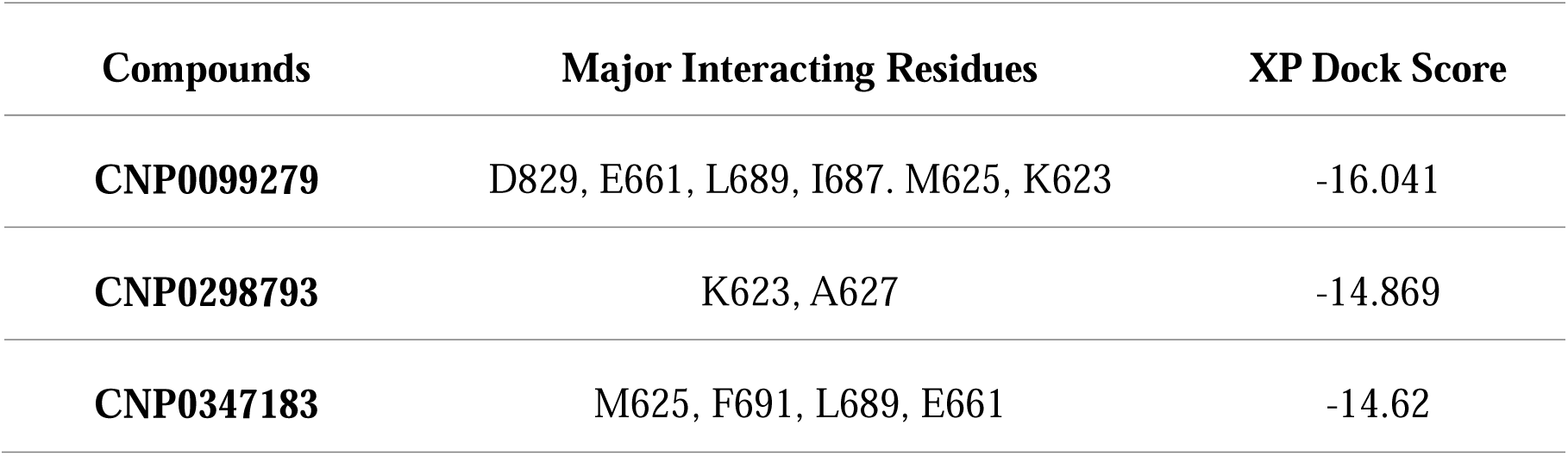
Top 3 compounds with COCONUT ID, interacting residues, and XP docking score (kcal/mol) of best-docked candidates, with zero golden triangle violations.

| Compounds | Major Interacting Residues | XP Dock Score |
| --- | --- | --- |
| **CNP0099279** | D829, E661, L689, I687, M625, K623 | -16.041 |
| **CNP0298793** | K623, A627 | -14.869 |
| **CNP0347183** | M625, F691, L689, E661 | -14.62 |

### 3.5. ADME/T Predictions

In this study, we utilized QikProp, SwissADME, ADMETlab 2.0, ProTox-II and pkCMS web servers to predict the ADMET properties of the best-performing compounds.

CNP0099279, CNP0298793, and CNP0347183 exhibit diverse molecular structures and pharmacokinetic profiles, indicative of their potential as drug candidates. CNP0099279 possesses a molecular weight of 439.180 g/mol, a significant dipole moment of 5.386, and 11 hydrogen bond donors and 12 acceptors. Its density of 1.068 g/mL suggests solid-state nature. CNP0298793, with a molecular weight of 498.531 g/mol, showcases a dipole moment of 9.852, 5 hydrogen bond donors, and 7 acceptors. CNP0347183, weighing 443.502 g/mol, presents a dipole moment of 5.437, 7 hydrogen bond donors, and 10 acceptors. Moreover, CNP0099279 has 10 rotatable bonds and a nine-atom ring system, while CNP0298793 and CNP0347183 have 1 and 13 rotatable bonds, respectively.

In terms of drug-likeness, CNP0099279 exhibits moderate water solubility (log mol/L = − 2.523) and skin permeability (log Kp = −2.735). CNP0298793 demonstrates moderate water solubility (log mol/L = −3.128) and skin permeability (log Kp = −2.735), with good Caco-2 permeability (0.956) and high human intestinal absorption (80.681%). CNP0347183 displays moderate water solubility (log mol/L = −2.873) and skin permeability (log Kp = −2.735), with relatively negative Caco-2 permeability (−0.474) and human intestinal absorption (27.248%).

Pharmacokinetically, CNP0099279 exhibits low plasma protein binding (16.916%) and a moderate volume of distribution (log L/kg = 0.406). CNP0298793 shows high plasma protein binding (97.46%), significant distribution throughout the body, and limited CNS penetration (logBB = −1.139). CNP0347183 displays low plasma protein binding (48.589%) and a moderate volume of distribution at steady state (log L/kg = −0.056), with low permeability across the blood-brain barrier (logBB = −0.8) and into the central nervous system (logPS = − 4.343). Additionally, CNP0298793 serves as a substrate for P-glycoprotein and inhibits its activity, suggesting the modulation of drug transport processes.

**Table 3.**
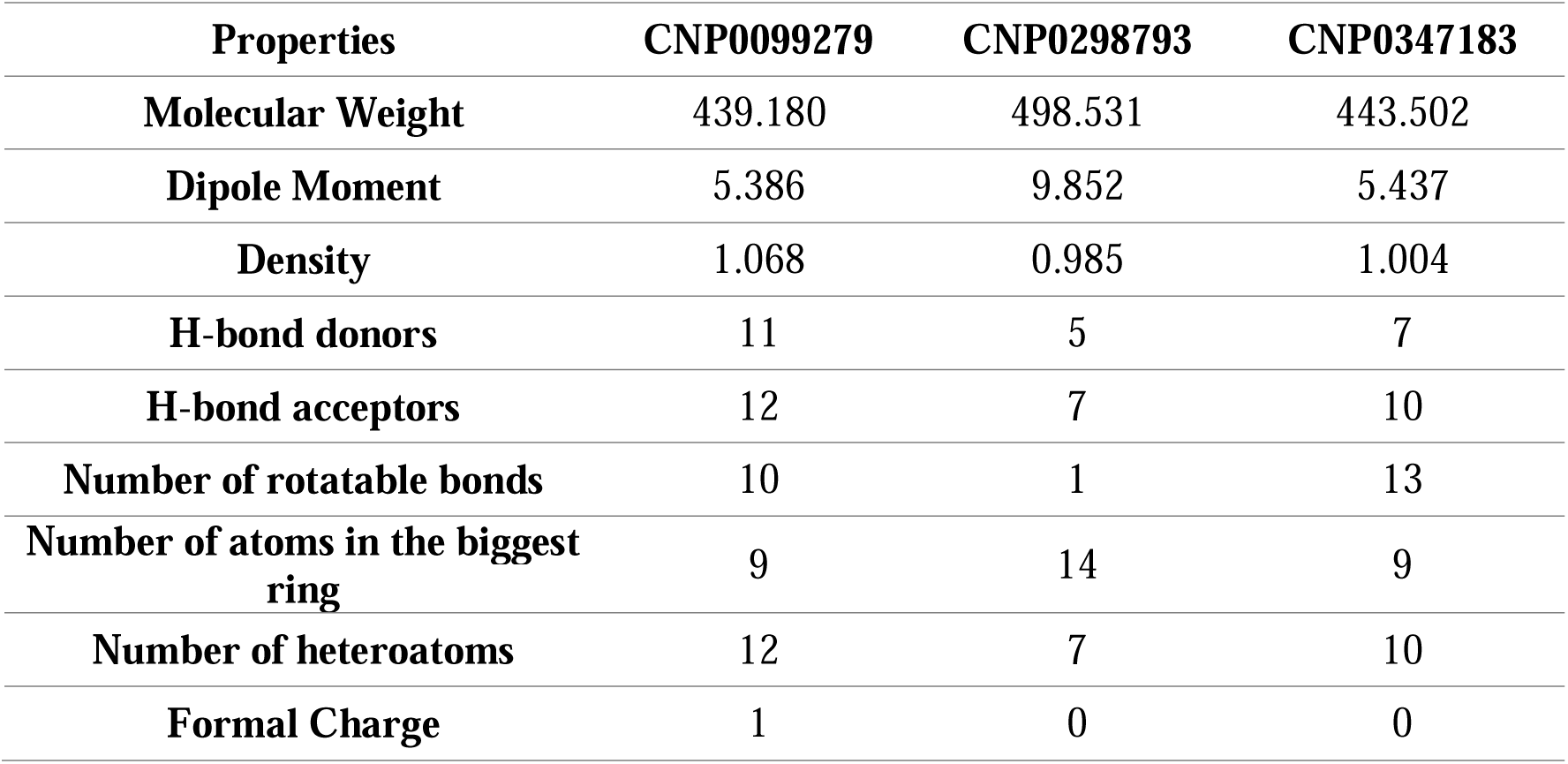
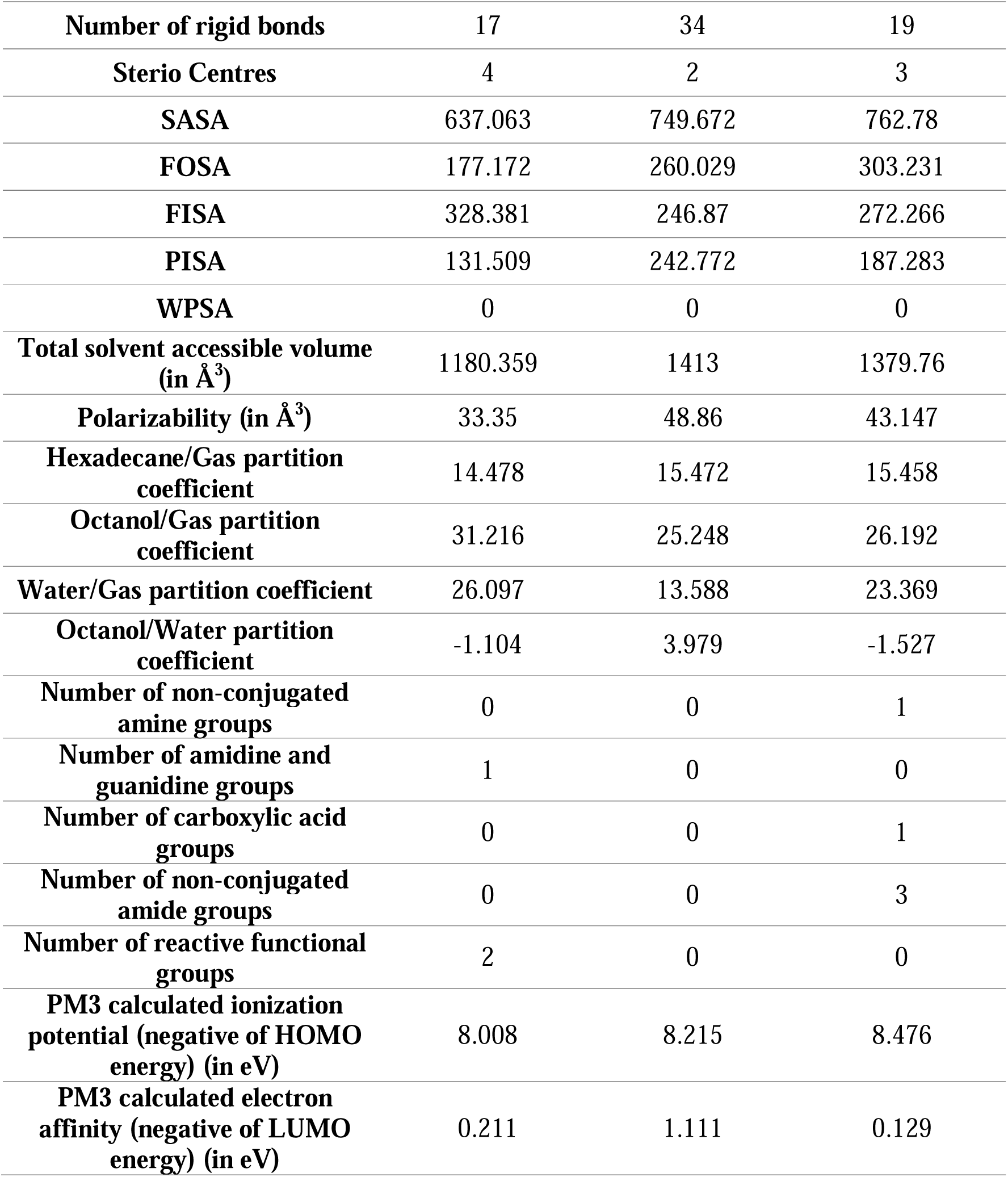
Physiochemical properties of the three screened compounds.

### 3.6. Molecular Dynamics Simulation

#### 3.6.1. RMSD analysis

Molecular Dynamics simulations of 100 ns time trajectory were conducted to study the molecular interactions that underlie the CNP0099279, CNP0298793, and CNP0347183 ligands binding in the protein pocket 6JQR. The MD simulation trajectories were initially composed of the RMSD analysis of each system. The RMSD value for 6JQR backbone unbound aligned with the RMSD values for 6JQR-CNP0099279, 6JQR-CNP0298793 and 6JQR-CNP0347183 complexes.

In Figure 6, it was observed the backbone atoms of 6JQR protein remained almost stable during the 100 ns simulation period for all the complexes A and C. For complex 6(B), the backbone of the protein increased in the beginning but stabilized from 35ns till the end of the simulation. We also observed the convergence in the RMSD values of the amino acid residue atoms of free 6JQR protein, indicating good trajectory stability in the complex. The RMSD values of 6JQR- CNP0099279 complex increased from 2.5Å at 0.15 ns to 3.39Å at 32.7ns and were then stabilized and averaged 3.25Å to end of MD trajectory, while the RMSD values of 6JQR- CNP0298793 complex went from 2.29Å for 1.7 ns to 3.47Å at 36.9 ns and thereafter these RMSD were not changed and stabilized with an average of 3.46Å till the end of the MD trajectory. The RMSD of the complex 6(C), were almost stable all throughout the simulation, with very minor fluctuations. The average values of its RMSD plot was 2.97 Å.

**Figure 6.**
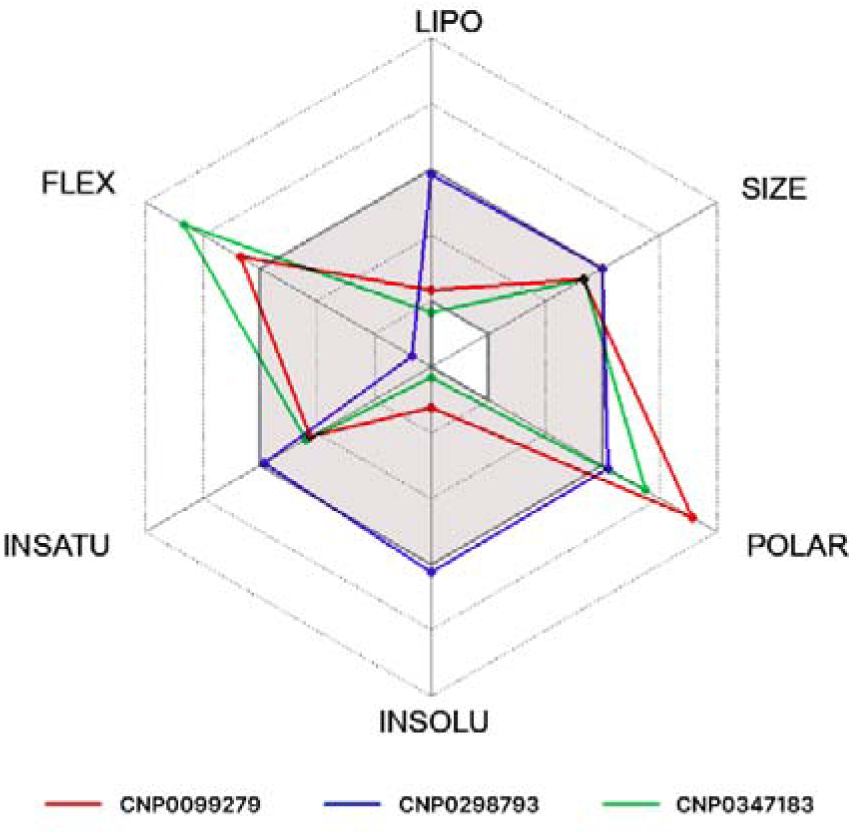
The bioavailability radar plot (obtained from SwissADME) depicting the excellent drug-likeliness of the two compounds (The reddish-brown area represents the optimal range for each property)

#### 3.6.2. RMSF Analysis

The RMSF computations for the backbone atoms of 6JQR analysed (see Figure 7). The average RMSF values of the protein backbone was under 2Å for all the three complexes. The average RMSF was found to be 1.51Å, 1.54Å, and 1.72Å respectively for the complexes with ligands CNP0099279, CNP0298793, and CNP0347183 respectively. Some slight fluctuations were seen at the residues interacting with the ligand atoms.

**Figure 7.**
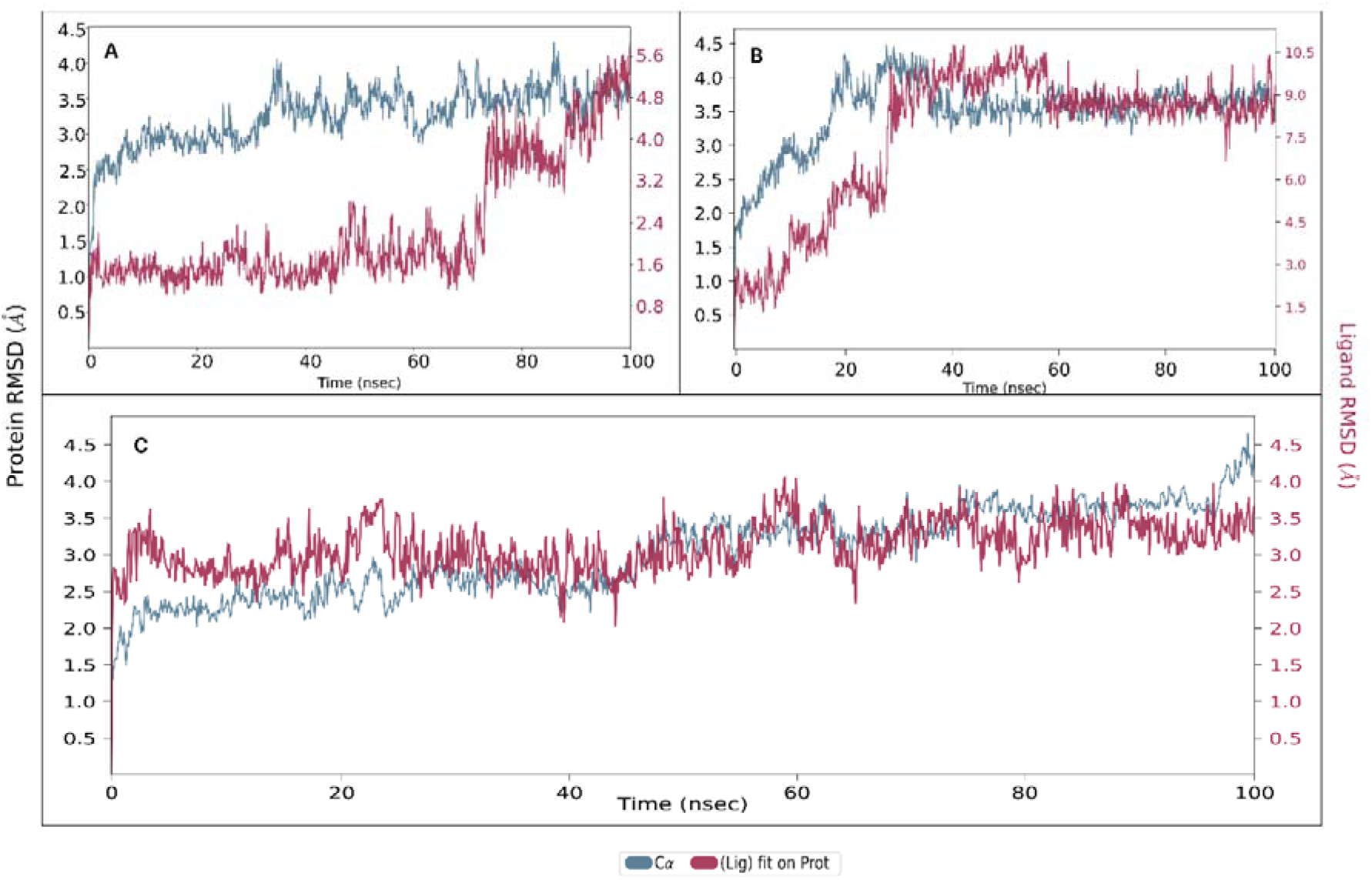
The line graph showing RMSD values of the complex structure extracted from ligand fit protein (ligand concerning protein) atoms. (A: *CNP0099279*; B: *CNP0298793; C: CNP0347183*)

RMSF values of the atoms of ligands CNP0099279, CNP0298793, and CNP0347183 were computed to describe the dynamic behaviour of the ligands within the 6JQR protein pocket (see Figure 8). The data indicate that some fluctuations in ligand CNP0099279 did not exceed 4.75Å for atom number 21, with an average of 2Å. For the ligands CNP0298793, and CNP0347183, the RMSF was maximum for atom number 34 and 25 with values of 4.94 Å and 2.65 Å respectively. The average RMSF values for these 2 ligands were 3.85 Å and 1.75 Å respectively. These slight fluctuations in ligand structures can be attributed to the flexible nature of the examined ligands. Nonetheless, the molecular structures of both ligands remained stable, with 6JQR backbone atoms exhibiting stability over 100 ns in aqueous conditions.

**Figure 8.**
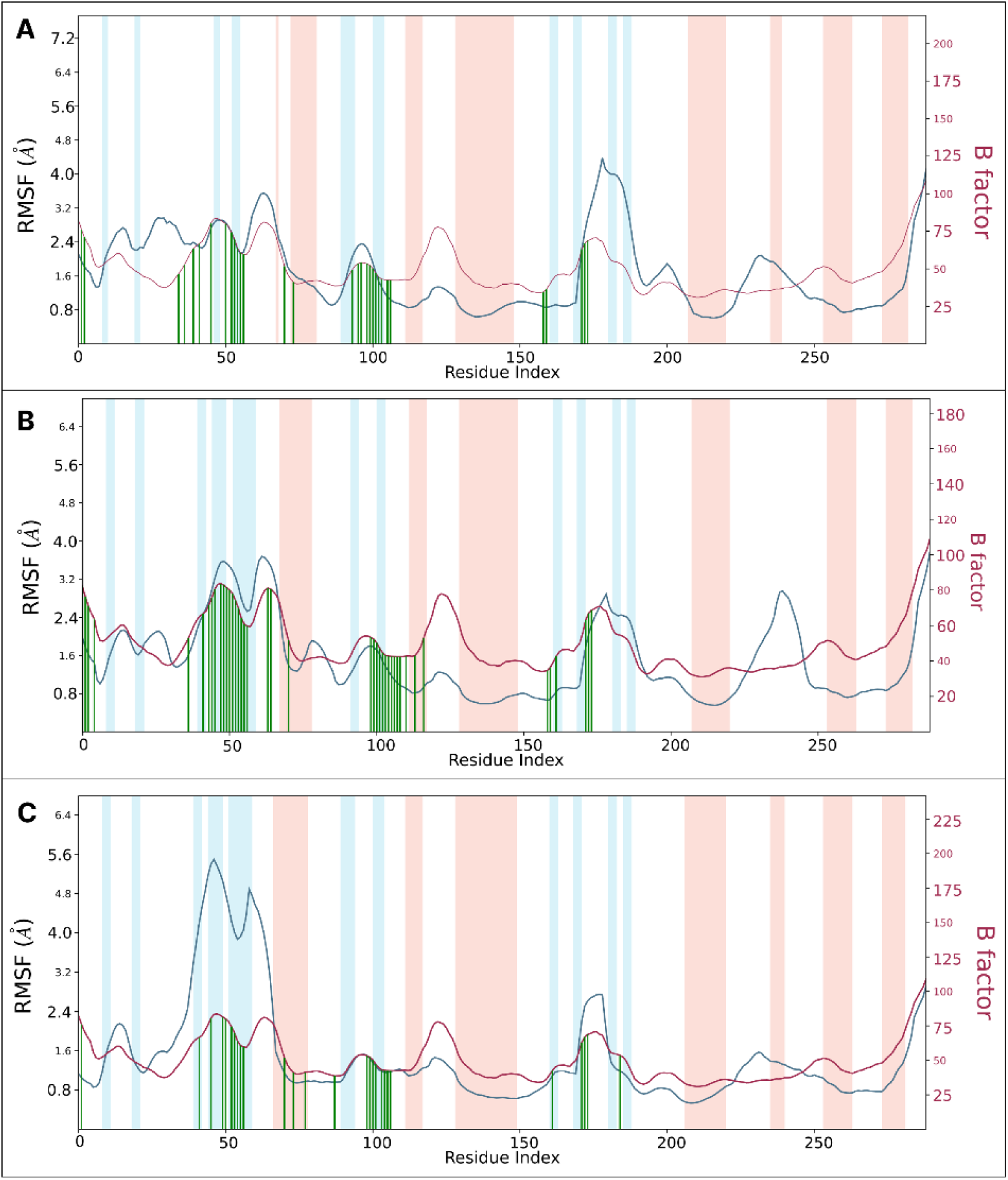
The line graph showing RMSF values of the complex structure extracted from protein residues backbone. The peaks of the blue line graph indicate the areas of the protein that fluctuate during the simulation. The maroon line graph indicates the B factor. The blue regions represent the alpha-helices and the orange region indicates the beta-helices. Protein residues that interact with the ligand are shown in green- coloured vertical lines.

**Figure 9.**
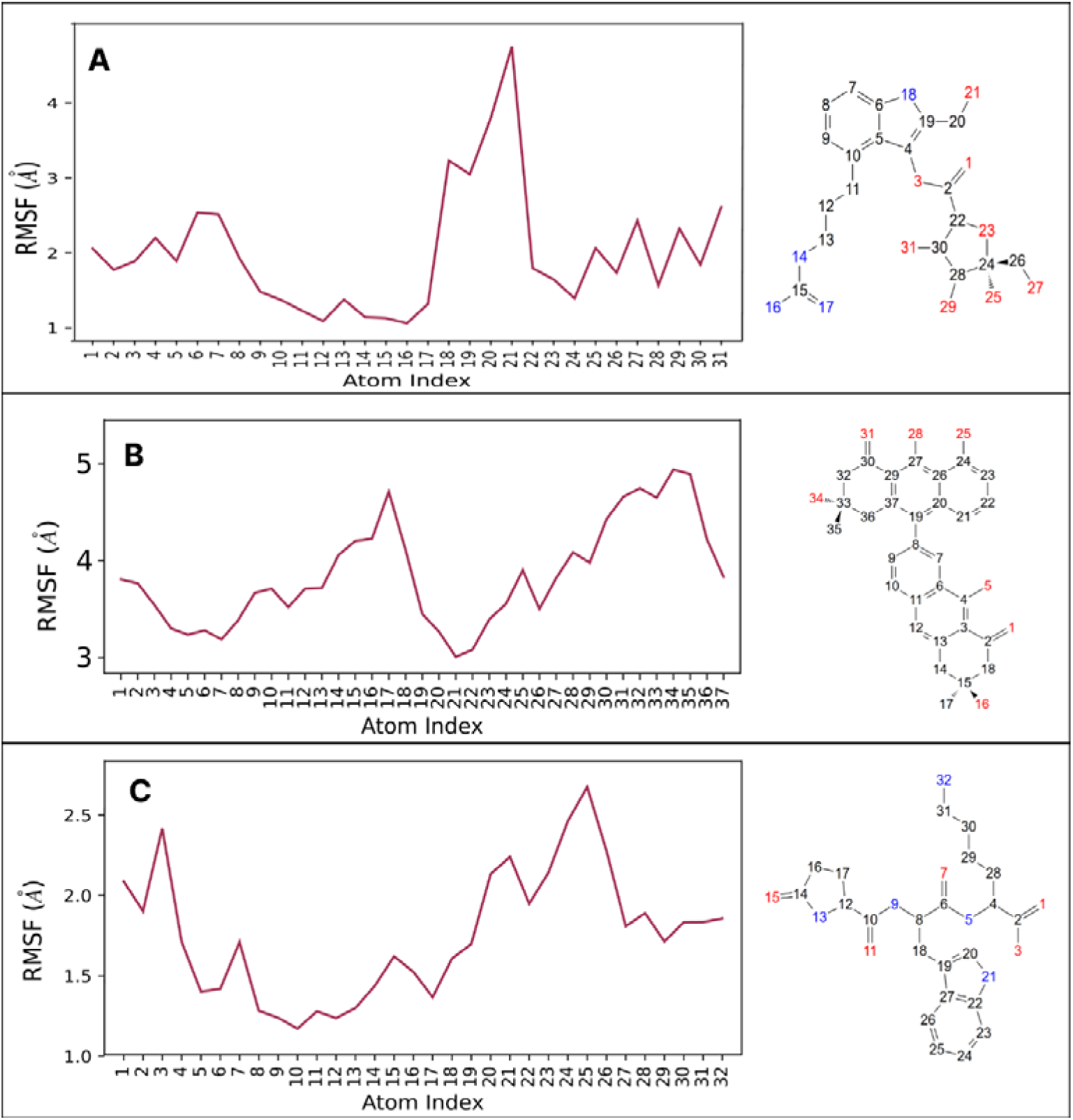
RMSF of Ligands’ atoms

#### 3.6.3. Protein-ligand contact dynamics

In the present study, it is identified four types of interactions within MD simulation: hydrogen bonds, hydrophobic interactions, water bridges, and ionic bonds. The timeline interactions of the residues of the protein with the ligands are shown in Figure S4, which shows a stable interaction.

The compound CNP0099279 showed significant hydrogen bond interactions with ASP829, GLU661, and LYS623 of our target protein that persisted throughout the 100 ns simulation. Furthermore, the compounds CNP0298793, and CNP0347183 showed extensive H-bond interactions with residues MET625, LEU689, ASP829, GLU661, PHE691 and, CYS695 and ARG815 respectively. CNP0099279 showed additional interactions with amino acids such as PHE691 and PHE621 majorly with H-bond and hydrophobic interactions. Compound CNP0298793 showed additional interactions with residues like LYS623 via water bridge and H-bonds, whereas CNP0347183 showed interactions with PHE621 (majorly hydrophobic), ASP698 (majorly water bridge) and, PHE830 (majorly hydrophobic and water bridge).

#### 3.6.4. Properties analysis of ligands CNP0099279, CNP0298793, and CNP0347183

To validate the high stability of the examined complexes in an aqueous environment, we assessed the dynamic properties of ligands CNP0099279, CNP0298793, and CNP0347183, which contribute to their stable interaction with the 6JQR protein active sites. Figure 10(A,B,C) summarizes the six examined properties: Ligand RMSD, Radius of Gyration (rGyr), Molecular Surface Area (MolSA), Intramolecular Hydrogen Bonds (intraHB), Solvent Accessible Surface Area (SASA), and Polar Surface Area (PSA).

**Figure 10.**
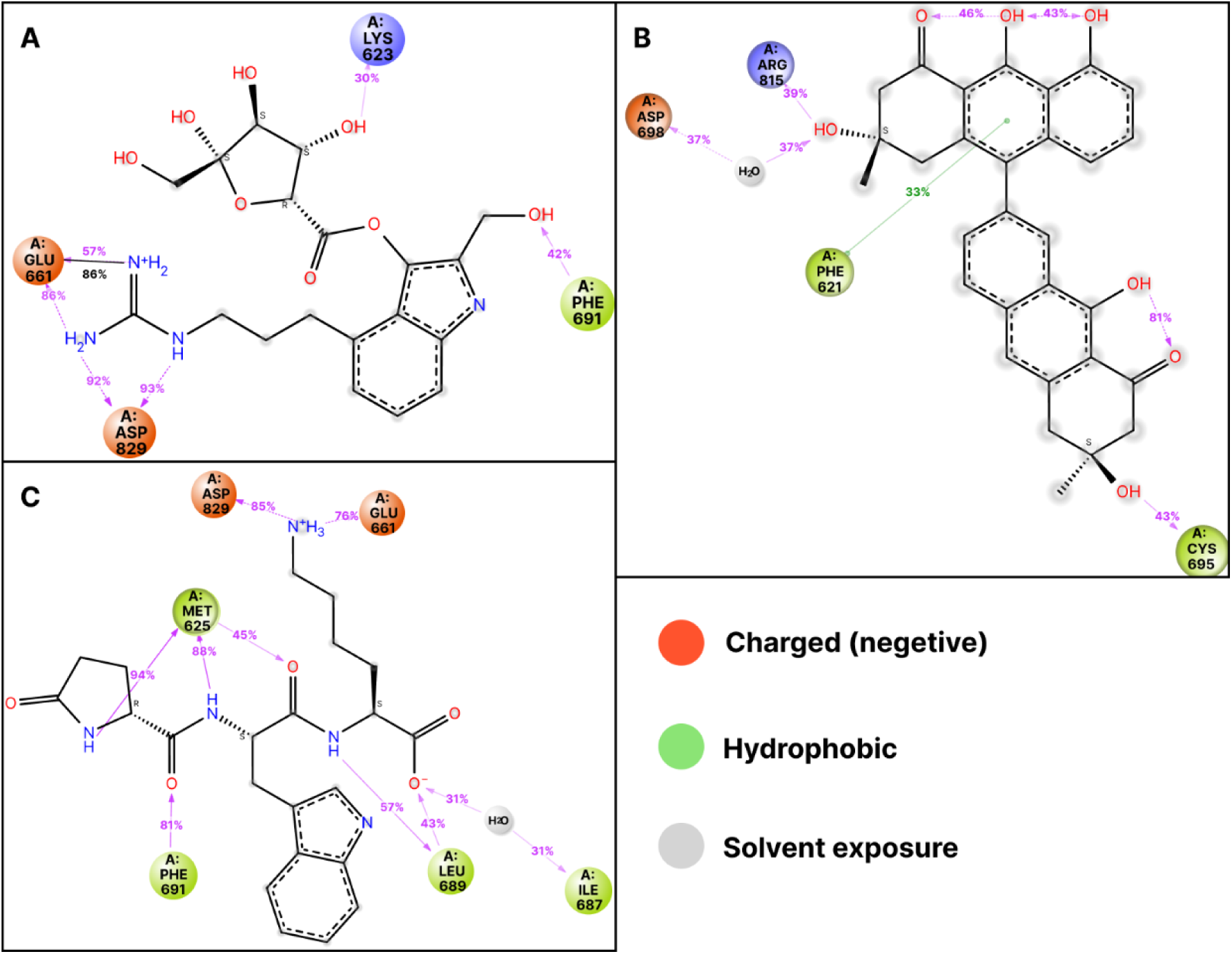
2D interaction diagram between protein-ligand complex (A) 6JQR-*CNP0099279* (B) 6JQR-*CNP0298793 (C) 6JQR- CNP0347183*

**Figure 11.**
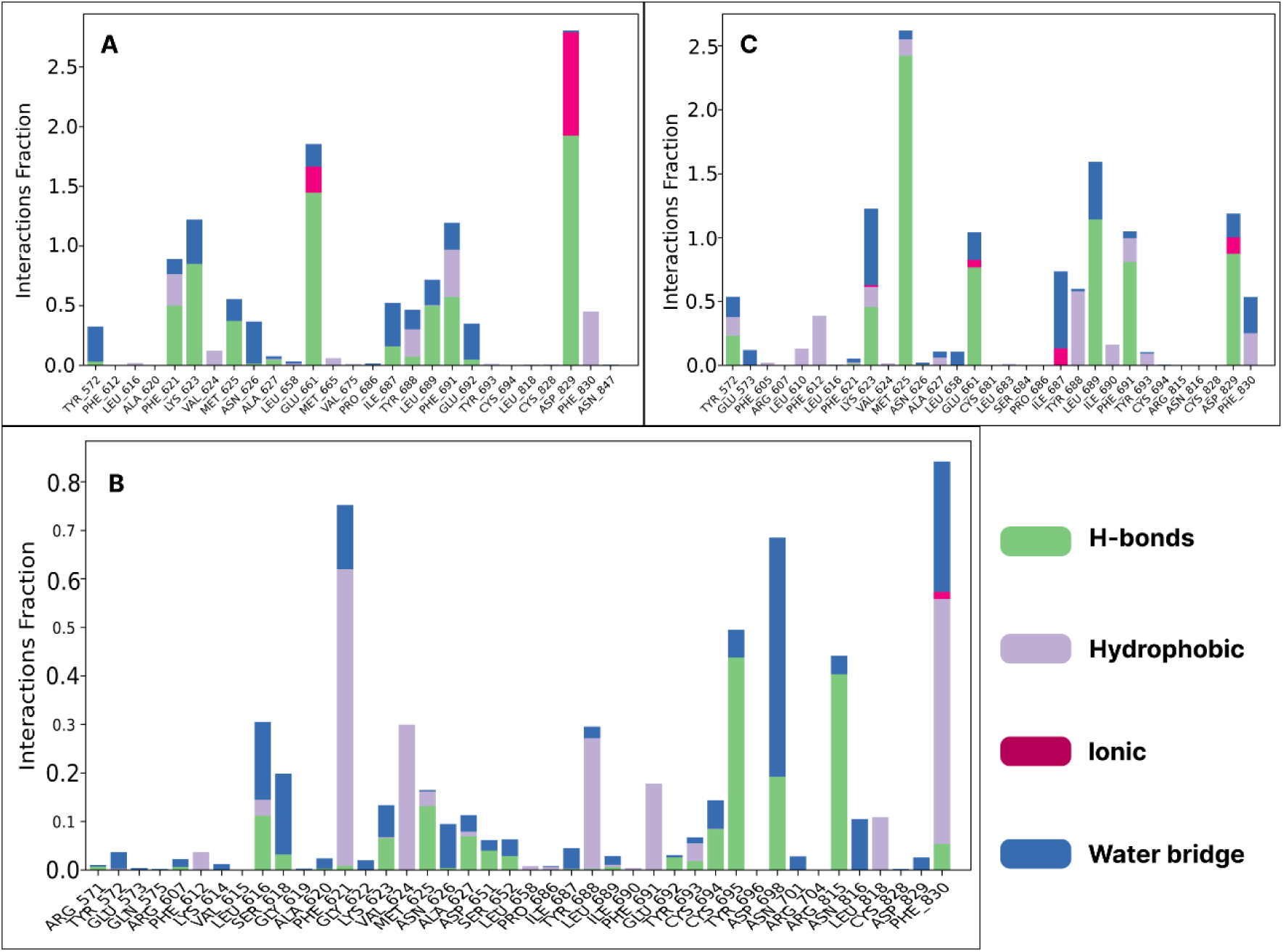
Interaction fraction histogram between protein-ligand complex (A) 6JQR-*CNP0099279* (B) 6JQR-*CNP0298793 (C) 6JQR-CNP0347183*

**Figure 12.**
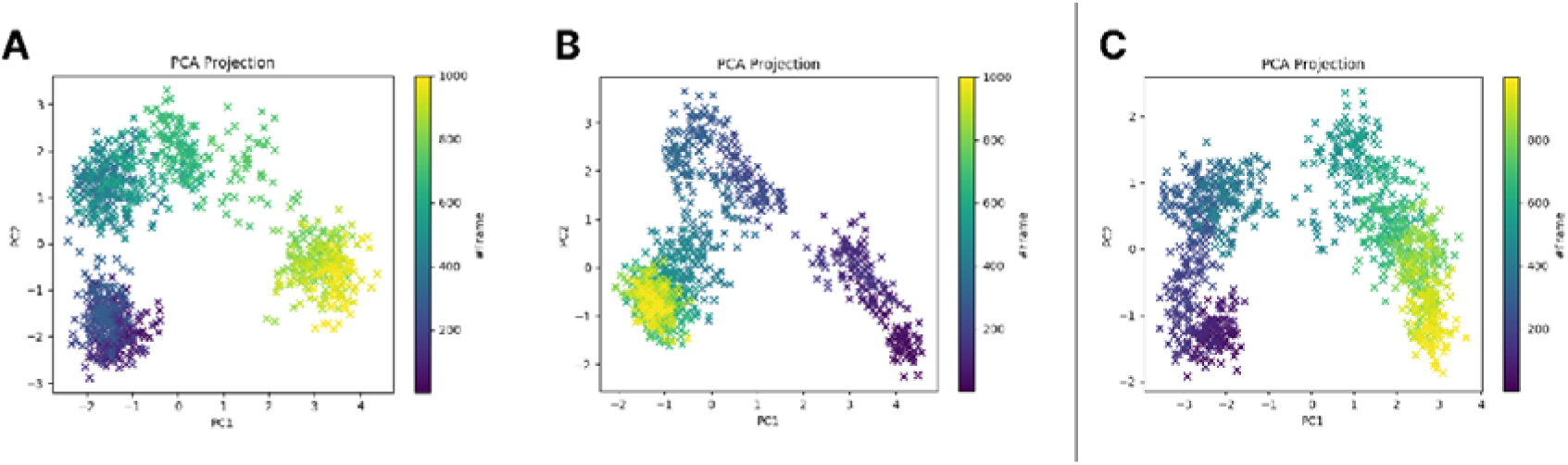
Standard 2D PCA scatter plot for the simulation trajectory of the complexes (A) 6JQR-*CNP0099279* (B) 6JQR-*CNP0298793 (C) 6JQR-CNP0347183*

**Figure 13.**
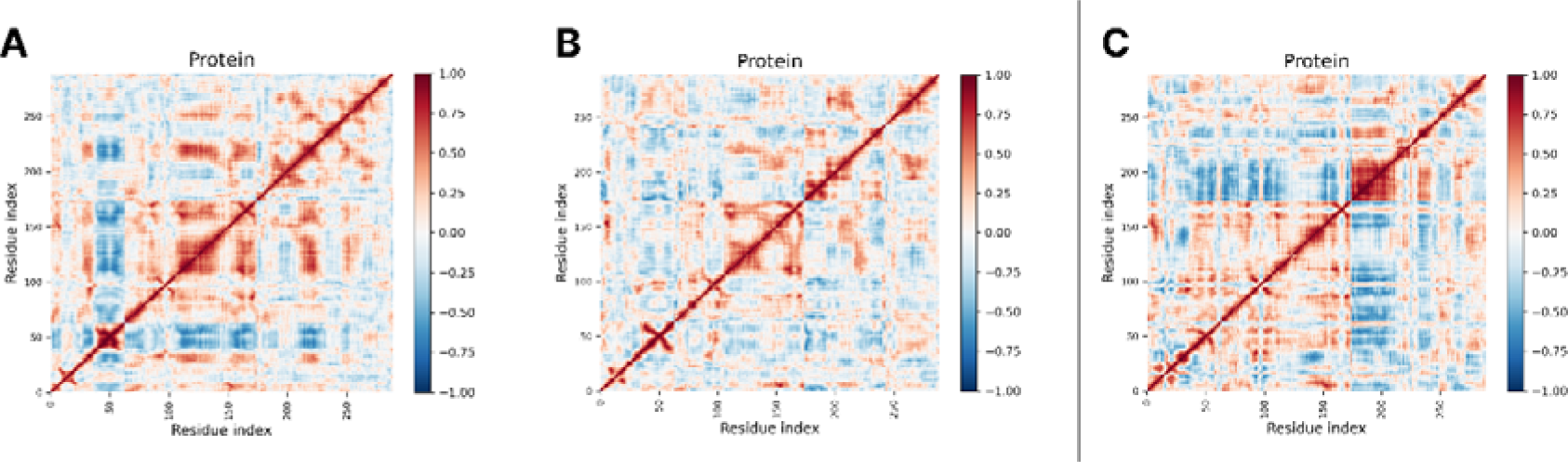
DCCM graph for the simulation trajectory of the complexes (A) 6JQR-*CNP0099279* (B) 6JQR-*CNP0298793 (C) 6JQR-CNP0347183*

From Figure S1(A,B,C), it is observed that the RMSD values of ligand CNP0099279, remained generally stable with an average value of 0.88Å till 73ns simulation time. After that, the RMSD showed a spike to an average of 2.53 Å, which was again almost stable. The RMSD values ranged between 0.44Å at 18.7 ns and 2.83Å at 89.6ns. On the other hand, the RMSD values of ligands CNP0298793, and CNP0347183 remained almost stable with an average of 0.76Å and 1.87 respectively throughout the 100 ns simulation, varying between 0.49Å at 54.4 ns and 1.18Å at 95.5ns for CNP0298793, whereas for CNP0347183 it is 1.6Å at 28ns and 2.9Å at 22ns. The slight variations in RMSD values for all the three ligands (<∼3Å) during the 100 ns MD simulation indicate the high stability of ligands ZINC000043204001 and ZINC000064540314 in the 6JQR protein pocket.

The average stability of rGyr values during the 100 ns MD simulation for ligands CNP0099279, CNP0298793, and CNP0347183 is 4.45Å, 5.25Å and 4.98Å, respectively. The intraHB, MolSA, SASA, and PSA plots in Figure S1(A, B, C) show good consistency in the dynamics of ligands CNP0099279, CNP0298793, and CNP0347183 within the 6JQR protein background. The presence of hydrogen bonds in the structures of ligands CNP0099279, CNP0298793, and CNP0347183, as indicated by the intraHB curves, could enhance the stability of these ligands in the 6JQR active pocket.

For the properties MolSA, SASA, and PSA of ligands CNP0099279, CNP0298793, and CNP0347183, values were observed in the range of (351-390Å, 415.3-429.5Å, and 394.8- 438.7Å), (21-218Å, 30.6-370.9Å, and 12.7-164Å), and (322-396Å, 231-261.6 Å, and 263.3-327.9Å), respectively. These findings further support the stability of ligands CNP0099279, CNP0298793, and CNP0347183 in the 6JQR protein environment.

## 4. Principal Component Analysis (PCA) and Dynamical Cross-Correlation Matrix (DCCM) Analysis

The PCA plot for CNP0099279 reveals three distinct clusters, indicating that the FLT3-ligand complex transitions between various conformational states during the simulation. The well-separated data points and clear color gradient show significant shifts in principal components (PC1 and PC2), suggesting that the complex spends time in stable or semi-stable conformations before transitioning. This smooth transition implies that CNP0099279 induces multiple stable conformations in the FLT3 protein. In contrast, the PCA plot for CNP0298793 shows a linear distribution of data points with less clustering, suggesting continuous conformational changes rather than distinct states. The narrow range on the PC2 axis and gradual color gradient imply that CNP0298793 induces a more flexible interaction with FLT3, with less defined stable conformations. The PCA plot for CNP0347183 displays two noticeable clusters with a continuous distribution between them, indicating at least two conformational states and fluid transitions. The clear trajectory between clusters suggests a balance of stability and flexibility, implying that CNP0347183 offers an intermediate scenario between the stability of CNP0099279 and the flexibility of CNP0298793.

CNP0099279 appears to be the most stable and likely the strongest FLT3 inhibitor candidate, inducing distinct conformational states in FLT3. CNP0347183 offers a balanced approach with some stability and flexibility, while CNP0298793 may need modifications to enhance its stability and specificity as an FLT3 inhibitor.

The DCCM graph for FLT3 with CNP0099279 displays a mix of positive (red) and negative (blue) correlations. Strong positive correlations along the diagonal indicate coordinated movement of neighboring residues, while off-diagonal clusters of positive and negative correlations around residues 50-100, 150-200, and 250-300 suggest that CNP0099279 stabilizes certain regions but allows flexibility in others. This indicates a balance between structural integrity and functional flexibility.

The DCCM graph for the FLT3 complex with CNP0298793 shows a more uniform pattern of positive correlations, particularly from residues 50 to 200. This suggests that CNP0298793 induces a more rigid and stable conformation in FLT3, enhancing structural integrity and reducing compensatory motions. The fewer and more localized negative correlations indicate a reduction in dynamic flexibility.

The DCCM graph for CNP0347183 reveals pronounced clusters of both positive and negative correlations. Strong positive correlations are observed in residues 0-50, 150-200, and 250- 300, with negative correlations in different regions. This indicates that CNP0347183 stabilizes specific domains while allowing flexibility in other areas, which may support functional responsiveness in FLT3.

The DCCM graphs illustrate that CNP0099279 balances stability and flexibility by stabilizing some regions while allowing others to be flexible. CNP0298793, on the other hand, induces a more uniformly stable structure with reduced flexibility. CNP0347183 stabilizes specific protein domains while promoting flexibility in others, potentially aiding in functional adaptability. These distinct patterns reveal how each ligand modulates FLT3’s dynamics, influencing its stability and functional interactions.

## 5. MM-GBSA Analysis

The parameters of free binding energies between the ligands were evaluated by the MM-GBSA simulations, which is the final step for checking the stability levels of the examined systems in the aqueous environment. The average values of calculated MM-GBSA energy parameters are presented in Table S4, containing binding free energies, Coulomb energy (Coulomb), Covalent bonding (Covalent), Hydrogen bonding (H-bond), lipophilic bonding (Lipo), π-π stacking interaction (Packing), solvent generalized bonding (SolvGB), and Van der Waals bonding energy (VDW) over the 100ns time frame.

From Table S6, the highest ΔGbind Total was shown by 6JQR-CNP0298793 Complex with average value of −80.14 kcal/mol and maximum value of −104.54 kcal/mol with a standard deviation (SD) of 10.45. The majority of it was contributed by ΔGbind Coulomb (−29.45 kcal/mol), ΔGbind Lipo (−24.92 kcal/mol) and, ΔGbind vdW (−56.33 kcal/mol). The next highest average ΔGbind Total was shown by the 6JQR-CNP0099279 complex and the 6JQR- CNP0298793 complex with values −73.75 kcal/mol (SD = 11.16) and −70.72 kcal/mol (SD = 11.43) respectively, as depicted in tables S4 and S5 respectively.

## 6. DISCUSSION

In this study, we employed *in silico* methods to discover novel FLT3 inhibitors for the treatment of acute myeloid leukemia (AML). The crux to our approach was the utilization of pharmacophore modelling, docking studies, ADMET analysis, molecular dynamics simulations, and MMGBSA calculations to assess the potential of identified compounds as drug candidates.

Pharmacophore modelling serves as a foundation in rational drug design by elucidating essential features necessary for ligand binding and activity at the target site (Choudhury and Sastry, 2019). We constructed a pharmacophore model based on a diverse set of known FLT3 inhibitors with confirmed pharmacological effects. This model consisted of distinct characteristics, including steric properties, hydrogen bond donor (HBD) and acceptor (HBA) groups, and electronic chemical attributes. The development of this model relied on the identification of active compounds with confirmed pharmacological effects targeting FLT3. However, it is crucial to note that the pharmacophore hypothesis match might have been set to a higher percentage if the number of diverse ligands was reduced from 250 to 100. This adjustment in the pharmacophore model parameters could potentially enhance its predictive capability and refine its ability to identify potent FLT3 inhibitors.

Docking studies were conducted to elucidate the interaction between identified compounds and FLT3. Our results revealed the interaction of the compounds with multiple novel residues of FLT3 like E661, L829, and M625, suggesting potential binding modes and highlighting possible residues for further *in vitro* validation. Notably, these interactions could represent novel ways of targeting FLT3 residues, which are crucial for the discovery of new inhibitors (Gokhale *et al*., 2019).

ADMET analysis was performed to assess the pharmacokinetic properties of the identified compounds. Among them, compound CNP0298793, which belongs to the super class of lignans, neolignans, and related compounds, exhibited significantly better ADMET properties. This compound falls under the class of Arylnaphthalene lignans, a subclass known for its antioxidant and anticancer activities. Lignans are secondary metabolites with diverse biological activities, especially in the context of anticancer properties. The lignan CNP0298793, obtained from the Mitishamba database, remains within acceptable pharmacokinetic ranges, suggesting its potential as a promising drug candidate with favorable profiles.

Molecular dynamics simulations provided valuable insights into the stability of protein-ligand complexes over time. Compounds CNP0298793, CNP0347183, and CNP0099279 demonstrated minimal fluctuations in protein root mean square deviation (RMSD) and root mean square fluctuation (RMSF), indicating stable interactions within the FLT3 active site. None of the RMSD values of all systems exceeded the 4Å threshold, suggesting a likely stability of these ligands inside the protein active pocket (Kufareva and Abagyan, 2012). Furthermore, the efficacy, selectivity, and side effects of a drug candidate depend on various factors, with protein-ligand interactions being of paramount importance. These interactions, including hydrogen bonds, hydrophobic bonds, water bridge bonds, and ionic bonds, contribute to the compound’s potency and selectivity, as highlighted by Fu *et al*. (2018). Strong interaction fractions (>1.5), particularly those involving residues such as Asp829 and Met625 with compound CNP0099279 and CNP0347183 respectively, having significant H-bond interactions, were observed indicating strong and stable binding to the protein, potentially inhibiting its activity. MMGBSA analysis reaffirmed the favourable binding energies of all three compounds, positioning them as promising drug candidates against FLT3, and presenting a comprehensive *in silico* approach for the discovery of novel FLT3 inhibitors in AML. The compound CNP0347183 exhibit the best binding energy over 100ns simulation.

The compound CNP0347183, derived from the GNPS (Global Natural Products Social Molecular Networking) database, is categorized under the super class of organic polymers and belongs to the class of polypeptides. Specifically, it is a pyroglutamyl-tryptophanyl- lysine tripeptide. Anti-cancer peptides (ACPs) like CNP0347183, composed of short sequences of amino acids, can inhibit tumor cell proliferation, migration, or suppress tumor angiogenesis, and are less likely to induce drug resistance. This compound exhibited the best binding energy over 100ns simulation, reaffirming its potential as a potent inhibitor of FLT3. CNP0099279, obtained from the Traditional Chinese Medicine Database (TCMDB@Taiwan), falls under the super class of organoheterocyclic compounds, with indoles as its direct parent class. The indole moiety is recognized for its exceptional role in anticancer drug development, particularly due to its ability to interact with a wide range of proteins and enzymes. The presence of a nitrogen atom within the indole structure allows for strong hydrogen bond formation with target proteins, enhancing its binding affinity and therapeutic potential.

The promising pharmacological properties of the positions the compounds for further investigation through *in vitro* and *in vivo* studies for potential clinical translation.

## 7. CONCLUSION

Our study employs a comprehensive computational framework to investigate novel FLT3 inhibitors sourced from natural compounds, leveraging the COCONUT database for structure-based drug discovery using pharmacophore modelling and screening, molecular docking, ADMET and subsequent molecular dynamics simulations. Through rigorous docking studies using HTVS, SP, and XP docking approaches, we identified potential inhibitors exhibiting high affinity towards FLT3. Among them, ligands CNP0099279, CNP0298793, and CNP0347183 emerged as top candidates with the highest docking scores, suggesting their strong binding affinity to FLT3.

To understand the molecular mechanisms underlying their inhibitory effects, we performed molecular dynamics simulations over a 100 ns trajectory. Our analysis revealed that ligands CNP0099279, CNP0298793, and CNP0347183 form stable complexes with FLT3, as evidenced by their consistent RMSD values throughout the simulation period. Notably, interactions with key residues such as Met625, Leu689, Glu661, and Asp829 were observed, highlighting their pivotal role in stabilizing the ligand-protein complexes.

Furthermore, our study underscores the importance of elucidating the detailed molecular mechanisms of these compounds in exerting AML tumor-suppressive effects. The identification of novel residues targeted by these inhibitors opens avenues for further exploration and validation.

Moving forward, it is imperative to conduct extensive *in vitro* and *in vivo* validation studies to confirm the efficacy and safety profiles of CNP0099279, CNP0298793, and CNP0347183. Additionally, detailed mechanistic studies are warranted to elucidate the precise mode of action and downstream signalling pathways affected by these inhibitors.

## Supporting information

Supplimentary_FLT3

