## Supplementary material for "Discovery of natural compounds as novel FMS-like tyrosine kinase-3 (FLT3) therapeutic inhibitors for the treatment of acute myeloid leukemia: an *in silico* approach": Supplimentary_FLT3

Table 1 Physiochemical properties of the three screened compounds

| Properties | CNP0099279 | CNP0298793 | CNP0347183 |
| --- | --- | --- | --- |
| Molecular Weight | 439.180 | 498.531 | 443.502 |
| Dipole Moment | 5.386 | 9.852 | 5.437 |
| Density | 1.068 | 0.985 | 1.004 |
| H-bond donors | 11 | 5 | 7 |
| H-bond acceptors | 12 | 7 | 10 |
| Number of rotatable bonds | 10 | 1 | 13 |
| Number of atoms in the biggest ring | 9 | 14 | 9 |
| Number of heteroatoms | 12 | 7 | 10 |
| Formal Charge | 1 | 0 | 0 |
| Number of rigid bonds | 17 | 34 | 19 |
| Sterio Centres | 4 | 2 | 3 |
| SASA | 637.063 | 749.672 | 762.78 |
| FOSA | 177.172 | 260.029 | 303.231 |
| FISA | 328.381 | 246.87 | 272.266 |
| PISA | 131.509 | 242.772 | 187.283 |
| WPSA | 0 | 0 | 0 |
| Total solvent accessible volume (in Å^3^) | 1180.359 | 1413 | 1379.76 |
| Polarizability (in Å^3^) | 33.35 | 48.86 | 43.147 |
| Hexadecane/Gas partition coefficient | 14.478 | 15.472 | 15.458 |
| Octanol/Gas partition coefficient | 31.216 | 25.248 | 26.192 |
| Water/Gas partition coefficient | 26.097 | 13.588 | 23.369 |
| Octanol/Water partition coefficient | -1.104 | 3.979 | -1.527 |
| Number of non-conjugated amine groups | 0 | 0 | 1 |
| Number of amidine and guanidine groups | 1 | 0 | 0 |
| Number of carboxylic acid groups | 0 | 0 | 1 |
| Number of non-conjugated amide groups | 0 | 0 | 3 |
| Number of reactive functional groups | 2 | 0 | 0 |
| PM3 calculated ionization potential (negative of HOMO energy) (in eV) | 8.008 | 8.215 | 8.476 |
| PM3 calculated electron affinity (negative of LUMO energy) (in eV) | 0.211 | 1.111 | 0.129 |

Table 2 Medicinal Chemistry properties of the three screened compounds

| Properties | CNP0099279 | CNP0298793 | CNP0347183 |
| --- | --- | --- | --- |
| Synthetic Accessibility Score | 4.760 | 4.056 | 4.059 |
| MCE-18 Value | 72.286 | 129.316 | 60.562 |
| PAINS | 0 alert | 0 alert | 0 alert |
| ALARM NMR Rule | 0 alert | 1 alert | 0 alert |
| BMS Rule | 0 alert | 0 alert | 0 alert |
| Chelator Rule | 0 alert | 0 alert | 0 alert |
| Lipinski’s Rule of 5 violations | 2 | 0 | 0 |
| Golden Triangle | Accepted | Accepted | Accepted |

Table 3 ADMET properties of the three screened compounds

| **Properties** | | **CNP0099279** | **CNP0298793** | **CNP0347183** |
| --- | --- | --- | --- | --- |
| **A** | Water solubility (log mol/L) | -2.523 | -3.128 | -2.873 |
|  | Caco-2 permeability (Papp in 10^-6^ cm/s) | -0.349 | 0.956 | -0.474 |
|  | Human intestinal absorption (in %) | 0.2 | 80.681 | 27.248 |
|  | Skin Permeability (log K_p_) | -2.735 | -2.735 | -2.735 |
|  | Human oral absorption (in %) | 10.35 | 79.859 | 0 |
|  | P-glycoprotein substrate | Yes | Yes | Yes |
|  | P-glycoprotein I inhibitor | No | Yes | No |
|  | P-glycoprotein II inhibitor | No | Yes | No |
|  | MDCK Permeability (in nm/sec) | 2.54 | 17.393 | 0.679 |
| **D** | Volume of distribution at steady state (in log L/kg) | 0.406 | -1.525 | -0.056 |
|  | Plasma protein binding (in %) | 16.916 | 97.46 | 48.589 |
|  | Fraction of unbound plasms | 0.731 | 0.26 | 0.62 |
|  | BBB Permeability (in logBB) | -1.527 | -1.139 | -0.8 |
|  | CNS permeability (in logPS) | -4.9 | -3.144 | -4.343 |
| **M** | CYP2D6 substrate | No | No | No |
|  | CYP3A4 substrate | No | Yes | Yes |
|  | CYPIA2 inhibitor | No | No | No |
|  | CYP2C19 inhibitor | No | No | No |
|  | CYP2C9 inhibitor | No | Yes | No |
|  | CYP2D6 inhibitor | No | No | No |
|  | CYP3A4 inhibitor | No | No | No |
| **E** | Total Clearance (in log ml/min/kg) | 1.34 | -0.875 | 1.478 |
|  | Renal OCT2 substrate | No | No | No |
| **T** | AMES toxicity | No | No | No |
|  | Max. tolerated dose (human) | 0.043 | 0.247 | 0.186 |
|  | hERG I inhibitor | No | No | No |
|  | hERG II inhibitor | No | Yes | No |
|  | Oral Rat Acute Toxicity (LD50) | 2.429 | 2.353 | 1.998 |
|  | Oral Rat Chronic Toxicity (LOAEL) | 4.464 | 2.543 | 2.114 |
|  | Hepatotoxicity | Yes | Yes | Yes |
|  | Skin Sensitisation | No | No | No |
|  | *T Pyriformis* toxicity (µg/L) | 0.285 | 0.285 | 0.285 |
|  | Minnow toxicity (log mM) | 5.217 | 0.156 | 3.324 |
|  | DILI | Passed | Low Risk | Passed |
|  | FDA Maximum Recommended Daily Dose | Passed | Low Risk | Low risk |
|  | Carcinogenicity | Passed | Passed | Passed |
|  | Eye corrosion | Passed | Passed | Passed |
|  | Eye irritation | Passed | Passed | Passed |
|  | Respiratory toxicity | Passed | Passed | Passed |

Table 4 Post simulation MM-GBSA based Binding free energy (ΔG Bind in kcal/mol) for the 6JQR-CNP0099279 Complex

| Time (in ns) | ΔG_bind_ Total | ΔG_bind_ Coulomb | ΔG_bind_ Covalent | ΔG_bind_ Hbond | ΔG_bind_ Lipo | ΔG_bind_ Packing | ΔG_bind_ Solv_GB | ΔG_bind_ vdW |
| --- | --- | --- | --- | --- | --- | --- | --- | --- |
| 1 | -78.26 | -92.13 | 5.18 | -6.04 | -18.33 | -1.81 | 84.03 | -49.15 |
| 2 | -86.93 | -97.89 | 6.84 | -7.80 | -20.62 | -1.95 | 79.73 | -45.23 |
| 3 | -83.38 | -101.71 | 9.11 | -6.65 | -20.47 | -2.00 | 81.06 | -42.72 |
| 4 | -91.18 | -101.05 | 7.80 | -6.80 | -22.91 | -2.16 | 86.62 | -52.69 |
| 5 | -86.67 | -106.73 | 10.31 | -7.04 | -21.88 | -2.06 | 94.86 | -54.14 |
| 6 | -81.24 | -106.67 | 7.48 | -6.78 | -20.80 | -1.91 | 94.52 | -47.08 |
| 7 | -80.14 | -97.04 | 11.74 | -7.04 | -20.28 | -2.18 | 83.29 | -48.64 |
| 8 | -91.10 | -96.82 | 7.26 | -6.84 | -22.79 | -2.10 | 81.58 | -51.40 |
| 9 | -88.16 | -99.59 | 12.23 | -6.88 | -21.87 | -1.68 | 85.28 | -55.65 |
| 10 | -85.69 | -111.35 | 12.60 | -6.90 | -22.92 | -1.91 | 96.73 | -51.94 |
| 11 | -92.77 | -94.84 | 8.20 | -7.28 | -23.46 | -1.97 | 79.91 | -53.34 |
| 12 | -93.95 | -103.01 | 7.94 | -6.97 | -23.04 | -2.26 | 86.14 | -52.75 |
| 13 | -81.27 | -94.50 | 9.72 | -6.88 | -22.08 | -1.53 | 83.41 | -49.40 |
| 14 | -80.68 | -90.99 | 11.25 | -6.59 | -22.23 | -1.03 | 81.42 | -52.51 |
| 15 | -87.05 | -104.97 | 9.79 | -7.05 | -21.82 | -2.71 | 94.35 | -54.65 |
| 16 | -80.77 | -86.20 | 10.66 | -6.98 | -19.10 | -1.07 | 76.84 | -54.91 |
| 17 | -78.02 | -97.70 | 8.17 | -6.32 | -21.50 | -1.29 | 94.75 | -54.12 |
| 18 | -90.55 | -95.02 | 9.91 | -7.10 | -23.42 | -2.28 | 81.68 | -54.32 |
| 19 | -90.61 | -93.66 | 7.38 | -6.86 | -22.65 | -1.68 | 79.99 | -53.14 |
| 20 | -91.97 | -115.32 | 5.62 | -7.60 | -22.79 | -1.90 | 94.89 | -44.87 |
| 21 | -86.94 | -94.78 | 6.93 | -6.77 | -21.40 | -2.10 | 81.29 | -50.11 |
| 22 | -75.65 | -105.61 | 8.47 | -6.37 | -19.70 | -1.72 | 101.84 | -52.56 |
| 23 | -73.08 | -88.60 | 9.29 | -5.38 | -20.77 | -1.15 | 82.25 | -48.73 |
| 24 | -86.32 | -102.97 | 11.07 | -7.76 | -21.60 | -1.29 | 86.39 | -50.17 |
| 25 | -85.69 | -111.91 | 11.16 | -7.65 | -20.41 | -1.61 | 84.23 | -39.50 |
| 26 | -82.72 | -89.61 | 6.07 | -5.90 | -20.43 | -0.75 | 69.73 | -41.84 |
| 27 | -76.96 | -95.66 | 11.79 | -6.32 | -19.95 | -0.71 | 80.94 | -47.05 |
| 28 | -83.77 | -90.50 | 8.59 | -6.67 | -18.57 | -0.20 | 71.04 | -47.47 |
| 29 | -86.19 | -102.67 | 4.68 | -7.17 | -19.60 | -1.35 | 82.54 | -42.64 |
| 30 | -88.24 | -102.28 | 7.61 | -7.37 | -21.71 | -1.06 | 81.20 | -44.63 |
| 31 | -78.64 | -99.39 | 6.30 | -6.84 | -17.92 | -1.89 | 82.13 | -41.03 |
| 32 | -76.68 | -77.20 | 5.23 | -5.73 | -20.88 | -3.67 | 70.82 | -45.26 |
| 33 | -78.21 | -79.75 | 8.57 | -5.66 | -21.42 | -2.72 | 74.28 | -51.50 |
| 34 | -71.22 | -78.29 | 8.05 | -4.54 | -23.22 | -4.17 | 74.41 | -43.46 |
| 35 | -67.88 | -72.64 | 6.20 | -5.17 | -20.05 | -1.62 | 72.60 | -47.19 |
| 36 | -85.25 | -78.83 | 6.97 | -4.90 | -25.34 | -4.98 | 76.79 | -54.94 |
| 37 | -81.27 | -87.23 | 8.40 | -5.87 | -24.74 | -4.60 | 78.33 | -45.56 |
| 38 | -70.53 | -90.91 | 11.07 | -6.17 | -20.92 | -2.29 | 87.98 | -49.29 |
| 39 | -81.09 | -81.83 | 8.15 | -5.63 | -23.63 | -3.68 | 77.82 | -52.29 |
| 40 | -82.72 | -84.31 | 9.06 | -5.44 | -24.08 | -2.42 | 75.11 | -50.65 |
| 41 | -84.49 | -75.83 | 5.03 | -5.80 | -25.12 | -4.95 | 76.66 | -54.48 |
| 42 | -87.72 | -68.36 | 5.81 | -5.87 | -24.60 | -3.75 | 63.32 | -54.27 |
| 43 | -86.16 | -89.02 | 5.63 | -6.12 | -23.30 | -4.79 | 79.23 | -47.80 |
| 44 | -75.19 | -90.04 | 8.07 | -5.14 | -22.53 | -2.37 | 80.17 | -43.35 |
| 45 | -75.46 | -78.17 | 7.37 | -6.10 | -20.15 | -2.20 | 69.67 | -45.89 |
| 46 | -64.55 | -78.35 | 7.30 | -4.36 | -18.50 | -1.49 | 70.20 | -39.35 |
| 47 | -66.87 | -74.09 | 10.38 | -5.51 | -21.20 | -1.55 | 68.92 | -43.82 |
| 48 | -70.84 | -82.14 | 9.39 | -5.80 | -18.66 | -2.54 | 62.03 | -33.12 |
| 49 | -63.33 | -69.10 | 6.70 | -4.76 | -16.82 | -2.02 | 60.96 | -38.29 |
| 50 | -77.29 | -72.17 | 8.61 | -5.13 | -21.11 | -3.86 | 59.71 | -43.34 |
| 51 | -59.07 | -68.32 | 4.34 | -3.71 | -18.15 | -2.54 | 67.58 | -38.28 |
| 52 | -61.68 | -62.86 | 10.40 | -3.77 | -23.55 | -4.48 | 68.64 | -46.05 |
| 53 | -70.82 | -88.89 | 7.89 | -4.24 | -21.44 | -3.57 | 82.45 | -43.02 |
| 54 | -64.76 | -75.77 | 11.69 | -4.73 | -20.63 | -4.04 | 72.89 | -44.16 |
| 55 | -75.45 | -76.68 | 8.65 | -5.19 | -22.65 | -4.01 | 68.59 | -44.16 |
| 56 | -66.62 | -70.42 | 6.63 | -3.89 | -23.19 | -4.09 | 70.54 | -42.21 |
| 57 | -70.27 | -73.15 | 6.47 | -4.59 | -20.28 | -2.88 | 68.73 | -44.57 |
| 58 | -84.06 | -74.53 | 4.10 | -5.33 | -25.02 | -5.37 | 72.90 | -50.82 |
| 59 | -89.50 | -77.70 | 7.10 | -5.17 | -24.68 | -5.07 | 69.67 | -53.66 |
| 60 | -81.58 | -86.33 | 7.14 | -5.17 | -24.34 | -4.46 | 76.64 | -45.06 |
| 61 | -79.98 | -74.44 | 8.18 | -5.82 | -22.52 | -3.83 | 65.29 | -46.84 |
| 62 | -82.86 | -85.94 | 5.03 | -5.20 | -23.88 | -3.30 | 76.00 | -45.57 |
| 63 | -74.25 | -63.45 | 6.03 | -4.92 | -21.18 | -3.20 | 63.30 | -50.84 |
| 64 | -64.15 | -77.19 | 2.72 | -4.22 | -16.85 | -1.90 | 71.90 | -38.61 |
| 65 | -69.00 | -67.84 | 6.77 | -5.01 | -18.53 | -3.66 | 66.18 | -46.91 |
| 66 | -74.19 | -78.71 | 8.77 | -4.84 | -23.28 | -4.96 | 74.03 | -45.21 |
| 67 | -70.80 | -67.96 | 5.94 | -4.72 | -20.04 | -4.27 | 68.66 | -48.40 |
| 68 | -64.98 | -84.02 | 8.46 | -4.11 | -19.15 | -3.22 | 78.99 | -41.92 |
| 69 | -62.43 | -80.93 | 10.46 | -4.45 | -20.64 | -3.89 | 81.42 | -44.40 |
| 70 | -68.64 | -92.04 | 9.06 | -5.02 | -18.13 | -1.61 | 79.63 | -40.52 |
| 71 | -66.75 | -75.23 | 4.98 | -5.06 | -16.41 | -1.77 | 70.60 | -43.86 |
| 72 | -60.66 | -78.79 | 4.16 | -3.71 | -14.82 | -0.30 | 70.88 | -38.07 |
| 73 | -61.12 | -75.72 | 2.98 | -4.76 | -16.37 | -0.93 | 70.63 | -36.96 |
| 74 | -51.52 | -94.98 | 6.13 | -4.42 | -11.64 | -0.22 | 85.41 | -31.80 |
| 75 | -63.02 | -86.23 | 5.54 | -4.84 | -15.72 | -1.97 | 79.62 | -39.42 |
| 76 | -70.85 | -103.28 | 7.40 | -5.58 | -15.25 | -0.77 | 84.71 | -38.07 |
| 77 | -66.37 | -87.56 | 7.65 | -5.53 | -14.69 | -0.80 | 70.24 | -35.67 |
| 78 | -62.69 | -83.96 | 7.33 | -3.95 | -17.34 | -1.63 | 77.96 | -41.09 |
| 79 | -52.26 | -91.79 | 8.59 | -5.00 | -12.29 | -0.08 | 78.86 | -30.53 |
| 80 | -69.90 | -92.58 | 5.50 | -6.13 | -14.82 | -1.63 | 74.76 | -35.01 |
| 81 | -61.80 | -94.38 | 7.32 | -5.60 | -13.10 | -0.44 | 75.41 | -31.01 |
| 82 | -55.79 | -96.75 | 9.35 | -4.53 | -16.39 | -2.01 | 88.02 | -33.48 |
| 83 | -68.60 | -107.55 | 5.76 | -5.14 | -16.93 | -1.17 | 88.73 | -32.30 |
| 84 | -71.15 | -97.06 | 5.46 | -5.34 | -17.40 | -1.85 | 83.17 | -38.13 |
| 85 | -70.47 | -99.14 | 5.78 | -5.09 | -16.77 | -3.61 | 87.29 | -38.92 |
| 86 | -70.89 | -98.73 | 8.51 | -5.38 | -16.73 | -1.78 | 80.30 | -37.08 |
| 87 | -68.61 | -84.56 | 4.51 | -4.83 | -17.94 | -1.73 | 74.18 | -38.24 |
| 88 | -69.43 | -84.01 | 6.88 | -4.75 | -18.62 | -1.61 | 69.24 | -36.57 |
| 89 | -65.66 | -76.04 | 7.23 | -4.96 | -17.28 | -1.77 | 65.55 | -38.39 |
| 90 | -56.64 | -82.31 | 8.58 | -4.98 | -13.40 | -3.12 | 69.80 | -31.20 |
| 91 | -62.90 | -87.05 | 5.02 | -5.41 | -14.70 | -1.31 | 72.59 | -32.03 |
| 92 | -49.17 | -86.09 | 7.53 | -4.53 | -13.09 | -1.05 | 75.92 | -27.85 |
| 93 | -57.02 | -79.01 | 7.55 | -4.98 | -18.25 | -2.32 | 70.09 | -30.09 |
| 94 | -59.94 | -85.48 | 7.54 | -4.98 | -14.71 | -1.92 | 71.24 | -31.63 |
| 95 | -63.70 | -102.74 | 6.09 | -4.88 | -16.29 | -1.65 | 82.99 | -27.21 |
| 96 | -66.72 | -92.19 | 6.77 | -5.18 | -14.79 | -2.34 | 71.13 | -30.12 |
| 97 | -50.64 | -75.59 | 7.53 | -4.83 | -12.00 | -1.26 | 66.11 | -30.61 |
| 98 | -57.53 | -81.45 | 7.07 | -5.37 | -14.30 | -1.41 | 70.48 | -32.55 |
| 99 | -57.23 | -90.79 | 6.36 | -4.86 | -14.28 | -1.99 | 79.74 | -31.41 |
| 100 | -59.20 | -80.25 | 10.29 | -5.13 | -16.34 | -3.44 | 66.77 | -31.11 |
| MAX | **-93.95** | **-115.32** | **2.72** | **-7.80** | **-25.34** | **-5.37** | **59.71** | **-55.65** |
| MIN | **-49.17** | **-62.86** | **12.60** | **-3.71** | **-11.64** | **-0.08** | **101.84** | **-27.21** |
| AVG | **-73.75** | **-87.58** | **7.64** | **-5.60** | **-19.64** | **-2.33** | **77.20** | **-43.44** |
| SD | **11.16** | **11.84** | **2.08** | **1.02** | **3.42** | **1.26** | **8.54** | **7.67** |

Table 5 Post simulation MM-GBSA based Binding free energy (ΔG Bind in kcal/mol) for the 6JQR-CNP0298793 Complex

| Time (in ns) | ΔG_bind_ Total | ΔG_bind_ Coulomb | ΔG_bind_ Covalent | ΔG_bind_ Hbond | ΔG_bind_ Lipo | ΔG_bind_ Packing | ΔG_bind_ Solv_GB | ΔG_bind_ vdW |
| --- | --- | --- | --- | --- | --- | --- | --- | --- |
| 1 | -96.81 | -34.15 | 3.65 | -2.56 | -27.76 | -4.05 | 29.96 | -61.90 |
| 2 | -86.24 | -30.05 | 1.04 | -2.20 | -26.30 | -2.59 | 34.19 | -60.33 |
| 3 | -102.10 | -31.13 | 2.18 | -2.30 | -29.62 | -4.57 | 31.22 | -67.88 |
| 4 | -85.45 | -27.57 | 1.46 | -1.93 | -25.52 | -3.14 | 35.48 | -64.23 |
| 5 | -93.31 | -27.96 | 1.26 | -1.54 | -27.84 | -3.29 | 33.42 | -67.36 |
| 6 | -78.58 | -25.64 | 2.92 | -1.95 | -23.95 | -3.23 | 31.50 | -58.22 |
| 7 | -89.64 | -32.65 | 2.98 | -2.16 | -25.59 | -4.70 | 40.01 | -67.53 |
| 8 | -72.65 | -21.64 | 2.27 | -1.91 | -21.03 | -3.55 | 34.19 | -60.97 |
| 9 | -74.77 | -30.48 | 2.02 | -1.75 | -21.89 | -4.75 | 35.66 | -53.58 |
| 10 | -82.50 | -33.72 | 1.18 | -2.38 | -23.06 | -4.24 | 34.73 | -55.02 |
| 11 | -89.38 | -21.39 | 3.84 | -1.47 | -28.68 | -9.87 | 26.25 | -58.04 |
| 12 | -82.11 | -30.03 | 2.29 | -2.12 | -22.02 | -6.45 | 29.19 | -52.98 |
| 13 | -76.86 | -21.44 | 2.41 | -0.89 | -25.85 | -6.15 | 29.88 | -54.81 |
| 14 | -77.46 | -26.22 | 2.38 | -2.03 | -22.86 | -5.18 | 29.29 | -52.84 |
| 15 | -78.24 | -25.24 | 6.18 | -1.21 | -24.24 | -7.62 | 31.37 | -57.48 |
| 16 | -79.44 | -23.49 | 4.31 | -1.84 | -23.76 | -7.53 | 32.01 | -59.15 |
| 17 | -75.71 | -23.82 | 2.49 | -1.03 | -25.34 | -4.60 | 25.31 | -48.73 |
| 18 | -70.72 | -19.63 | 2.58 | -1.16 | -23.66 | -5.19 | 25.76 | -49.41 |
| 19 | -62.83 | -18.34 | 1.93 | -0.91 | -21.93 | -6.12 | 26.48 | -43.95 |
| 20 | -57.36 | -13.69 | 0.43 | 0.00 | -22.78 | -4.23 | 27.34 | -44.43 |
| 21 | -48.13 | -10.61 | 1.80 | -1.15 | -19.13 | -3.57 | 23.63 | -39.10 |
| 22 | -55.51 | -19.15 | 1.69 | -0.59 | -20.53 | -3.95 | 27.19 | -40.17 |
| 23 | -49.62 | -7.02 | 2.14 | -0.92 | -18.45 | -4.75 | 21.31 | -41.94 |
| 24 | -50.43 | -6.04 | 1.83 | -1.39 | -19.57 | -4.31 | 21.23 | -42.18 |
| 25 | -45.62 | -10.20 | 2.75 | -0.01 | -16.91 | -3.91 | 18.79 | -36.13 |
| 26 | -60.29 | -17.30 | 2.32 | -0.75 | -19.90 | -3.47 | 22.84 | -44.03 |
| 27 | -63.10 | -14.88 | 10.15 | -1.27 | -23.31 | -4.12 | 22.14 | -51.81 |
| 28 | -75.94 | -16.16 | 5.47 | -1.68 | -26.56 | -4.89 | 17.92 | -50.04 |
| 29 | -52.13 | -10.24 | 3.56 | -0.57 | -19.70 | -5.01 | 17.08 | -37.26 |
| 30 | -44.23 | -9.16 | 2.20 | -0.52 | -17.57 | -1.90 | 16.99 | -34.27 |
| 31 | -68.13 | -28.19 | 4.29 | -1.12 | -22.48 | -4.43 | 26.02 | -42.21 |
| 32 | -57.31 | -19.08 | 6.85 | -1.48 | -20.82 | -3.62 | 21.44 | -40.59 |
| 33 | -69.58 | -21.10 | 5.57 | -1.63 | -23.14 | -4.12 | 24.06 | -49.22 |
| 34 | -65.66 | -17.67 | 1.97 | -1.06 | -20.64 | -4.94 | 20.76 | -44.08 |
| 35 | -60.63 | -12.75 | 2.97 | -0.79 | -17.49 | -3.31 | 21.20 | -50.46 |
| 36 | -57.74 | -15.96 | 3.31 | -0.55 | -18.89 | -1.80 | 24.99 | -48.83 |
| 37 | -49.87 | -10.01 | 1.16 | -0.59 | -17.68 | -1.43 | 23.76 | -45.10 |
| 38 | -67.94 | -14.09 | 0.63 | -1.50 | -21.93 | -5.88 | 20.10 | -45.26 |
| 39 | -63.95 | -10.88 | 0.09 | -0.86 | -22.57 | -6.62 | 22.19 | -45.28 |
| 40 | -59.18 | -15.37 | 1.58 | -1.25 | -19.85 | -6.97 | 23.32 | -40.65 |
| 41 | -76.06 | -22.75 | -3.83 | -1.41 | -22.55 | -7.32 | 26.27 | -44.47 |
| 42 | -65.38 | -10.31 | 1.54 | -0.98 | -24.09 | -5.78 | 22.45 | -48.21 |
| 43 | -70.73 | -15.21 | 2.59 | -0.79 | -24.84 | -5.35 | 22.86 | -49.98 |
| 44 | -63.80 | -12.76 | 1.31 | -1.04 | -22.11 | -4.59 | 24.42 | -49.02 |
| 45 | -57.54 | -7.94 | 0.91 | -0.41 | -22.25 | -2.14 | 23.74 | -49.45 |
| 46 | -52.57 | -10.63 | 0.58 | -0.23 | -20.19 | -1.72 | 23.66 | -44.04 |
| 47 | -69.37 | -20.54 | 1.73 | -1.67 | -21.96 | -4.22 | 23.73 | -46.45 |
| 48 | -60.10 | -16.62 | 1.96 | -1.41 | -19.11 | -4.44 | 19.14 | -39.63 |
| 49 | -55.69 | -15.63 | 1.68 | -1.40 | -17.73 | -4.75 | 22.96 | -40.82 |
| 50 | -68.42 | -21.66 | 0.81 | -1.43 | -20.22 | -4.87 | 22.07 | -43.12 |
| 51 | -57.33 | -21.26 | 1.45 | -1.36 | -17.69 | -4.00 | 25.69 | -40.16 |
| 52 | -51.26 | -19.14 | 2.72 | -1.75 | -14.95 | -2.86 | 22.61 | -37.89 |
| 53 | -59.43 | -11.50 | 1.72 | -0.89 | -22.33 | -4.57 | 21.44 | -43.30 |
| 54 | -66.09 | -21.99 | 0.85 | -1.44 | -20.91 | -3.53 | 24.82 | -43.89 |
| 55 | -76.83 | -18.49 | 2.62 | -0.78 | -29.03 | -3.54 | 26.77 | -54.39 |
| 56 | -65.03 | -14.83 | 3.98 | -0.59 | -25.26 | -2.14 | 25.82 | -52.01 |
| 57 | -68.33 | -23.12 | 0.32 | -0.62 | -23.50 | -5.56 | 28.44 | -44.30 |
| 58 | -69.47 | -20.30 | 1.75 | -1.05 | -24.42 | -5.40 | 27.84 | -47.88 |
| 59 | -69.39 | -20.71 | 3.02 | -1.17 | -24.59 | -7.46 | 27.82 | -46.29 |
| 60 | -73.33 | -18.09 | 2.41 | -1.07 | -25.25 | -5.26 | 23.65 | -49.71 |
| 61 | -70.49 | -24.35 | 2.07 | -1.15 | -22.39 | -3.36 | 25.64 | -46.95 |
| 62 | -75.45 | -22.89 | 8.15 | -1.29 | -26.39 | -6.43 | 26.85 | -53.45 |
| 63 | -72.09 | -23.91 | 3.98 | -1.23 | -23.72 | -7.67 | 28.93 | -48.47 |
| 64 | -83.37 | -19.27 | 2.86 | -1.08 | -29.69 | -8.21 | 24.96 | -52.94 |
| 65 | -75.53 | -22.12 | 0.81 | -1.09 | -25.31 | -3.61 | 25.97 | -50.19 |
| 66 | -74.50 | -20.01 | 3.58 | -1.05 | -27.31 | -6.36 | 26.71 | -50.06 |
| 67 | -69.55 | -16.57 | 3.36 | -1.06 | -23.30 | -5.49 | 21.58 | -48.07 |
| 68 | -85.12 | -20.18 | 1.79 | -1.09 | -28.95 | -6.89 | 23.87 | -53.67 |
| 69 | -77.64 | -14.88 | 2.32 | -1.01 | -26.56 | -5.72 | 22.72 | -54.51 |
| 70 | -82.13 | -21.00 | 2.86 | -1.22 | -27.57 | -7.23 | 26.02 | -53.98 |
| 71 | -81.63 | -19.83 | 1.89 | -1.07 | -26.96 | -6.62 | 23.84 | -52.88 |
| 72 | -67.07 | -21.67 | 1.72 | -1.08 | -22.25 | -3.46 | 28.67 | -49.00 |
| 73 | -73.81 | -24.21 | 1.78 | -1.15 | -23.84 | -4.07 | 28.95 | -51.28 |
| 74 | -70.19 | -19.60 | 5.25 | -1.11 | -22.93 | -4.59 | 24.44 | -51.65 |
| 75 | -85.13 | -19.63 | 2.33 | -1.09 | -28.25 | -6.64 | 23.79 | -55.64 |
| 76 | -80.31 | -19.48 | 2.19 | -1.08 | -26.93 | -6.00 | 24.96 | -53.96 |
| 77 | -83.61 | -28.69 | 2.83 | -1.07 | -25.97 | -6.34 | 27.82 | -52.19 |
| 78 | -80.18 | -31.93 | 4.30 | -1.08 | -26.26 | -6.80 | 35.25 | -53.66 |
| 79 | -71.04 | -9.10 | 2.48 | -0.60 | -27.12 | -7.73 | 24.59 | -53.55 |
| 80 | -82.47 | -18.36 | 2.01 | -1.10 | -26.90 | -5.84 | 23.63 | -55.89 |
| 81 | -81.06 | -19.71 | 3.44 | -1.07 | -27.11 | -7.91 | 23.16 | -51.86 |
| 82 | -67.95 | -16.13 | 1.73 | -1.15 | -22.88 | -6.57 | 23.34 | -46.28 |
| 83 | -73.80 | -24.97 | 2.31 | -1.08 | -26.24 | -6.33 | 29.47 | -46.95 |
| 84 | -78.84 | -23.31 | 2.02 | -1.06 | -26.67 | -7.47 | 26.61 | -48.95 |
| 85 | -71.78 | -18.71 | 2.59 | -1.09 | -24.69 | -6.47 | 25.05 | -48.45 |
| 86 | -76.50 | -23.96 | 2.33 | -1.06 | -26.27 | -3.85 | 26.25 | -49.95 |
| 87 | -71.42 | -18.66 | 3.04 | -1.07 | -24.34 | -5.75 | 22.63 | -47.27 |
| 88 | -81.60 | -30.91 | 3.44 | -1.07 | -27.29 | -7.03 | 30.40 | -49.14 |
| 89 | -79.38 | -26.43 | 3.99 | -1.11 | -25.57 | -5.46 | 26.90 | -51.69 |
| 90 | -78.37 | -20.67 | 2.94 | -1.08 | -26.51 | -5.90 | 23.50 | -50.64 |
| 91 | -70.92 | -12.33 | 2.30 | -0.55 | -24.61 | -6.70 | 21.51 | -50.54 |
| 92 | -62.56 | -7.31 | 0.92 | -0.05 | -26.78 | -8.36 | 25.33 | -46.32 |
| 93 | -73.23 | -21.76 | 2.17 | -1.06 | -23.73 | -5.83 | 25.31 | -48.34 |
| 94 | -77.00 | -24.90 | 2.88 | -1.26 | -25.30 | -7.04 | 27.48 | -48.85 |
| 95 | -61.59 | -16.05 | 4.51 | -1.36 | -20.95 | -3.43 | 23.02 | -47.33 |
| 96 | -61.20 | -12.38 | 1.79 | -0.87 | -22.34 | -7.62 | 27.07 | -46.84 |
| 97 | -64.65 | -12.59 | 2.29 | -0.58 | -22.69 | -4.57 | 22.81 | -49.32 |
| 98 | -84.44 | -26.19 | 1.39 | -1.06 | -28.02 | -7.51 | 27.88 | -50.93 |
| 99 | -68.37 | -18.77 | 1.56 | -1.34 | -23.85 | -7.31 | 25.43 | -44.10 |
| 100 | -72.65 | -22.41 | 9.32 | -1.05 | -27.56 | -5.59 | 27.00 | -52.36 |
| MAX | **-102.10** | **-34.15** | **-3.83** | **-2.56** | **-29.69** | **-9.87** | **16.99** | **-67.88** |
| MIN | **-44.23** | **-6.04** | **10.15** | **0.00** | **-14.95** | **-1.43** | **40.01** | **-34.27** |
| AVG | **-70.72** | **-19.63** | **2.58** | **-1.16** | **-23.66** | **-5.19** | **25.76** | **-49.41** |
| SD | **11.43** | **6.45** | **1.81** | **0.48** | **3.21** | **1.71** | **4.22** | **6.56** |

Table 6 Post simulation MM-GBSA based Binding free energy (ΔG Bind in kcal/mol) for the 6JQR-CNP0347183 Complex

| Time (in ns) | ΔG_bind_ Total | ΔG_bind_ Coulomb | ΔG_bind_ Covalent | ΔG_bind_ Hbond | ΔG_bind_ Lipo | ΔG_bind_ Packing | ΔG_bind_ Solv_GB | ΔG_bind_ vdW |
| --- | --- | --- | --- | --- | --- | --- | --- | --- |
| 1 | -75.34 | -26.18 | 7.86 | -3.61 | -23.69 | -4.73 | 27.77 | -52.76 |
| 2 | -72.19 | -23.73 | -0.78 | -3.85 | -23.68 | -4.31 | 38.01 | -53.85 |
| 3 | -54.58 | -27.96 | 5.14 | -3.49 | -19.83 | -3.82 | 43.85 | -48.46 |
| 4 | -57.72 | -23.08 | 1.88 | -3.53 | -20.37 | -2.62 | 35.78 | -45.79 |
| 5 | -81.82 | -36.65 | 4.50 | -4.26 | -26.42 | -3.27 | 40.65 | -56.38 |
| 6 | -93.03 | -47.36 | 4.92 | -5.87 | -28.80 | -3.57 | 42.20 | -54.55 |
| 7 | -87.60 | -38.11 | 2.78 | -5.81 | -25.72 | -1.70 | 35.61 | -54.65 |
| 8 | -88.27 | -38.66 | 8.10 | -6.16 | -27.41 | -3.82 | 35.94 | -56.26 |
| 9 | -95.42 | -32.06 | 3.48 | -5.58 | -28.38 | -3.95 | 29.36 | -58.29 |
| 10 | -82.90 | -29.82 | 7.15 | -5.03 | -27.59 | -3.04 | 33.15 | -57.71 |
| 11 | -88.68 | -37.56 | 3.98 | -6.02 | -26.47 | -3.49 | 35.24 | -54.35 |
| 12 | -86.91 | -34.01 | 5.32 | -5.59 | -24.83 | -2.14 | 29.70 | -55.37 |
| 13 | -85.18 | -31.25 | 6.03 | -5.56 | -26.22 | -4.21 | 34.25 | -58.21 |
| 14 | -94.96 | -43.78 | 5.56 | -5.46 | -25.26 | -3.21 | 36.20 | -59.01 |
| 15 | -84.96 | -48.74 | 5.50 | -5.89 | -26.25 | -3.18 | 40.70 | -47.10 |
| 16 | -82.87 | -36.34 | 5.09 | -5.53 | -24.23 | -2.95 | 33.61 | -52.52 |
| 17 | -91.58 | -35.46 | 4.65 | -6.25 | -25.60 | -4.94 | 38.73 | -62.70 |
| 18 | -73.87 | -28.36 | 3.28 | -4.70 | -20.98 | -1.00 | 31.32 | -53.43 |
| 19 | -85.21 | -43.99 | 5.47 | -4.85 | -23.03 | -1.57 | 38.44 | -55.68 |
| 20 | -75.74 | -33.87 | 7.36 | -4.87 | -21.94 | -1.12 | 31.84 | -53.14 |
| 21 | -65.98 | -13.82 | 4.49 | -3.05 | -23.14 | -2.34 | 25.25 | -53.38 |
| 22 | -84.03 | -24.80 | 8.64 | -4.35 | -25.51 | -1.37 | 21.74 | -58.37 |
| 23 | -72.79 | -23.04 | 6.28 | -4.65 | -21.95 | -0.71 | 24.20 | -52.91 |
| 24 | -77.77 | -19.29 | 8.46 | -3.16 | -26.31 | -3.60 | 26.99 | -60.85 |
| 25 | -78.80 | -30.53 | 4.42 | -3.30 | -23.88 | -0.92 | 32.33 | -56.92 |
| 26 | -81.27 | -32.37 | 11.84 | -3.97 | -27.15 | -1.50 | 35.52 | -63.63 |
| 27 | -92.93 | -36.93 | 9.67 | -4.05 | -29.69 | -4.83 | 32.95 | -60.05 |
| 28 | -89.91 | -35.92 | 6.35 | -3.64 | -26.39 | -4.33 | 33.72 | -59.71 |
| 29 | -80.26 | -29.84 | 8.26 | -4.30 | -22.74 | -4.42 | 33.90 | -61.12 |
| 30 | -86.04 | -36.10 | 4.18 | -4.12 | -25.74 | -2.93 | 34.56 | -55.90 |
| 31 | -91.25 | -36.81 | 8.39 | -4.02 | -28.87 | -3.95 | 33.33 | -59.32 |
| 32 | -90.08 | -44.09 | 5.61 | -3.99 | -25.07 | -1.91 | 38.84 | -59.46 |
| 33 | -82.99 | -37.02 | 6.59 | -4.14 | -25.23 | -2.08 | 33.34 | -54.44 |
| 34 | -90.21 | -32.83 | 6.69 | -4.38 | -28.71 | -4.38 | 27.53 | -54.13 |
| 35 | -95.12 | -37.25 | 9.05 | -4.36 | -30.48 | -3.76 | 33.17 | -61.49 |
| 36 | -87.41 | -34.45 | 5.45 | -3.81 | -27.86 | -3.02 | 29.95 | -53.68 |
| 37 | -85.05 | -31.14 | 3.53 | -4.28 | -25.22 | -1.36 | 30.34 | -56.92 |
| 38 | -85.53 | -37.55 | 4.27 | -4.00 | -26.10 | -5.22 | 36.80 | -53.75 |
| 39 | -81.06 | -42.13 | 5.72 | -4.33 | -24.60 | -1.88 | 38.25 | -52.09 |
| 40 | -92.25 | -46.03 | 8.99 | -4.20 | -25.46 | -0.55 | 33.64 | -58.64 |
| 41 | -84.95 | -38.37 | 9.84 | -4.27 | -28.09 | -1.17 | 35.49 | -58.38 |
| 42 | -90.82 | -32.57 | 5.06 | -4.13 | -25.60 | -1.32 | 29.06 | -61.31 |
| 43 | -75.75 | -41.11 | 8.17 | -4.45 | -21.14 | -1.16 | 36.44 | -52.50 |
| 44 | -83.42 | -25.82 | 6.91 | -5.09 | -27.19 | -0.13 | 26.99 | -59.10 |
| 45 | -81.58 | -20.28 | 3.79 | -4.91 | -25.85 | -1.44 | 23.79 | -56.68 |
| 46 | -86.40 | -16.21 | 4.94 | -5.08 | -29.54 | -0.65 | 21.67 | -61.54 |
| 47 | -81.67 | -32.16 | 4.92 | -5.45 | -25.05 | -1.82 | 26.67 | -48.78 |
| 48 | -83.12 | -26.81 | 4.28 | -4.99 | -26.36 | -2.36 | 28.14 | -55.03 |
| 49 | -85.96 | -29.56 | 6.72 | -5.11 | -25.38 | -1.13 | 26.70 | -58.20 |
| 50 | -92.74 | -30.22 | 4.95 | -4.80 | -26.47 | -1.50 | 28.33 | -63.03 |
| 51 | -82.44 | -12.57 | 9.98 | -4.33 | -27.19 | -0.40 | 19.74 | -67.68 |
| 52 | -83.68 | -27.73 | 4.74 | -5.11 | -27.68 | -0.64 | 30.59 | -57.86 |
| 53 | -84.92 | -30.00 | 8.49 | -4.81 | -26.68 | -0.01 | 28.01 | -59.93 |
| 54 | -94.27 | -35.54 | 9.24 | -5.22 | -31.35 | -1.89 | 28.23 | -57.74 |
| 55 | -86.25 | -20.49 | 7.60 | -4.34 | -25.76 | -0.25 | 18.64 | -61.65 |
| 56 | -89.19 | -30.90 | 4.48 | -5.35 | -24.44 | -1.10 | 29.15 | -61.03 |
| 57 | -90.67 | -30.36 | 4.89 | -4.51 | -28.93 | -3.48 | 32.52 | -60.81 |
| 58 | -87.13 | -28.12 | 9.62 | -5.40 | -30.21 | -2.08 | 27.69 | -58.63 |
| 59 | -104.53 | -35.09 | 6.14 | -5.10 | -31.59 | -3.57 | 28.97 | -64.28 |
| 60 | -99.64 | -26.57 | 3.79 | -4.98 | -33.55 | -3.69 | 28.57 | -63.21 |
| 61 | -100.45 | -27.40 | 5.66 | -4.84 | -30.56 | -2.10 | 24.07 | -65.28 |
| 62 | -86.52 | -18.09 | 2.37 | -3.63 | -28.05 | -1.40 | 20.42 | -58.15 |
| 63 | -96.28 | -39.92 | 5.73 | -4.53 | -29.39 | -2.88 | 41.44 | -66.75 |
| 64 | -80.29 | -40.90 | 4.01 | -4.42 | -27.39 | -1.79 | 49.63 | -59.42 |
| 65 | -83.08 | -33.21 | 2.81 | -4.68 | -27.95 | -1.80 | 39.79 | -58.05 |
| 66 | -79.54 | -34.07 | 5.64 | -4.66 | -28.85 | -1.27 | 39.81 | -56.13 |
| 67 | -82.27 | -38.90 | 3.26 | -4.59 | -28.03 | -2.30 | 49.94 | -61.65 |
| 68 | -77.07 | -40.22 | 3.71 | -4.64 | -24.42 | -0.44 | 49.57 | -60.64 |
| 69 | -80.93 | -33.12 | 6.31 | -4.12 | -29.90 | -2.39 | 45.57 | -63.29 |
| 70 | -75.53 | -31.14 | 5.86 | -3.99 | -25.36 | -2.61 | 41.19 | -59.47 |
| 71 | -71.86 | -38.33 | 5.08 | -4.42 | -23.28 | -1.24 | 45.87 | -55.55 |
| 72 | -69.12 | -35.86 | 4.53 | -4.48 | -20.93 | -1.45 | 41.74 | -52.67 |
| 73 | -84.55 | -38.45 | 0.31 | -4.83 | -26.21 | -0.20 | 41.43 | -56.60 |
| 74 | -67.55 | -19.55 | 4.53 | -3.35 | -21.43 | -0.27 | 30.00 | -57.49 |
| 75 | -67.02 | -18.38 | 4.60 | -3.31 | -22.39 | -0.41 | 26.88 | -54.02 |
| 76 | -64.92 | -26.80 | 2.65 | -4.43 | -21.05 | -0.77 | 34.95 | -49.48 |
| 77 | -69.51 | -28.56 | 2.15 | -3.87 | -19.91 | 0.00 | 35.74 | -55.07 |
| 78 | -75.60 | -33.56 | 0.47 | -4.30 | -21.46 | -0.70 | 33.76 | -49.82 |
| 79 | -74.65 | -35.89 | 3.41 | -3.16 | -20.83 | -0.88 | 35.89 | -53.18 |
| 80 | -79.25 | -31.78 | 1.38 | -3.87 | -23.62 | -0.40 | 35.38 | -56.35 |
| 81 | -79.70 | -27.05 | 2.81 | -4.60 | -25.09 | -0.86 | 31.27 | -56.17 |
| 82 | -67.42 | -37.14 | -0.50 | -5.17 | -21.94 | -0.77 | 47.78 | -49.69 |
| 83 | -77.16 | -42.85 | 2.54 | -5.52 | -22.88 | -0.56 | 46.48 | -54.37 |
| 84 | -71.65 | -31.26 | 2.38 | -3.73 | -21.70 | -1.56 | 35.50 | -51.27 |
| 85 | -69.15 | -38.43 | 2.00 | -5.12 | -20.67 | -0.60 | 45.05 | -51.38 |
| 86 | -76.64 | -39.32 | 2.35 | -5.09 | -23.51 | -1.99 | 40.91 | -49.99 |
| 87 | -72.82 | -26.04 | 3.54 | -2.85 | -23.67 | -0.48 | 33.32 | -56.64 |
| 88 | -73.82 | -10.00 | 1.65 | -2.07 | -23.75 | -1.18 | 14.55 | -53.02 |
| 89 | -66.03 | -13.77 | 4.32 | -2.64 | -20.56 | 0.00 | 19.73 | -53.11 |
| 90 | -67.06 | -24.54 | 3.70 | -2.60 | -19.69 | -0.09 | 29.35 | -53.19 |
| 91 | -66.96 | -16.48 | 3.86 | -3.09 | -20.56 | -0.15 | 25.24 | -55.78 |
| 92 | -66.35 | -10.97 | 3.32 | -3.07 | -19.51 | -0.43 | 17.03 | -52.72 |
| 93 | -50.87 | -14.05 | 1.89 | -2.98 | -13.40 | 0.00 | 21.57 | -43.89 |
| 94 | -69.82 | -14.73 | 3.35 | -2.71 | -23.06 | -0.64 | 21.58 | -53.61 |
| 95 | -67.55 | -10.02 | 4.72 | -3.31 | -22.55 | -0.85 | 18.50 | -54.04 |
| 96 | -67.61 | -9.49 | 2.67 | -2.69 | -21.55 | -0.01 | 20.89 | -57.44 |
| 97 | -65.85 | -14.85 | 2.53 | -2.46 | -19.84 | 0.00 | 23.03 | -54.26 |
| 98 | -64.92 | -7.71 | 3.54 | -2.53 | -19.76 | 0.00 | 14.51 | -52.96 |
| 99 | -66.87 | -9.82 | 3.16 | -2.52 | -23.04 | -0.33 | 20.48 | -54.80 |
| 100 | -61.34 | -2.69 | 1.73 | -2.23 | -19.52 | 0.00 | 12.50 | -51.13 |
| MAX | **-104.54** | **-48.74** | **-0.78** | **-6.25** | **-33.55** | **-5.22** | **12.50** | **-67.68** |
| MIN | **-50.87** | **-2.69** | **11.84** | **-2.07** | **-13.40** | **0.00** | **49.94** | **-43.89** |
| AVG | **-80.14** | **-29.75** | **4.93** | **-4.31** | **-24.92** | **-1.83** | **32.06** | **-56.33** |
| SD | **10.45** | **9.94** | **2.44** | **0.95** | **3.40** | **1.43** | **8.24** | **4.45** |

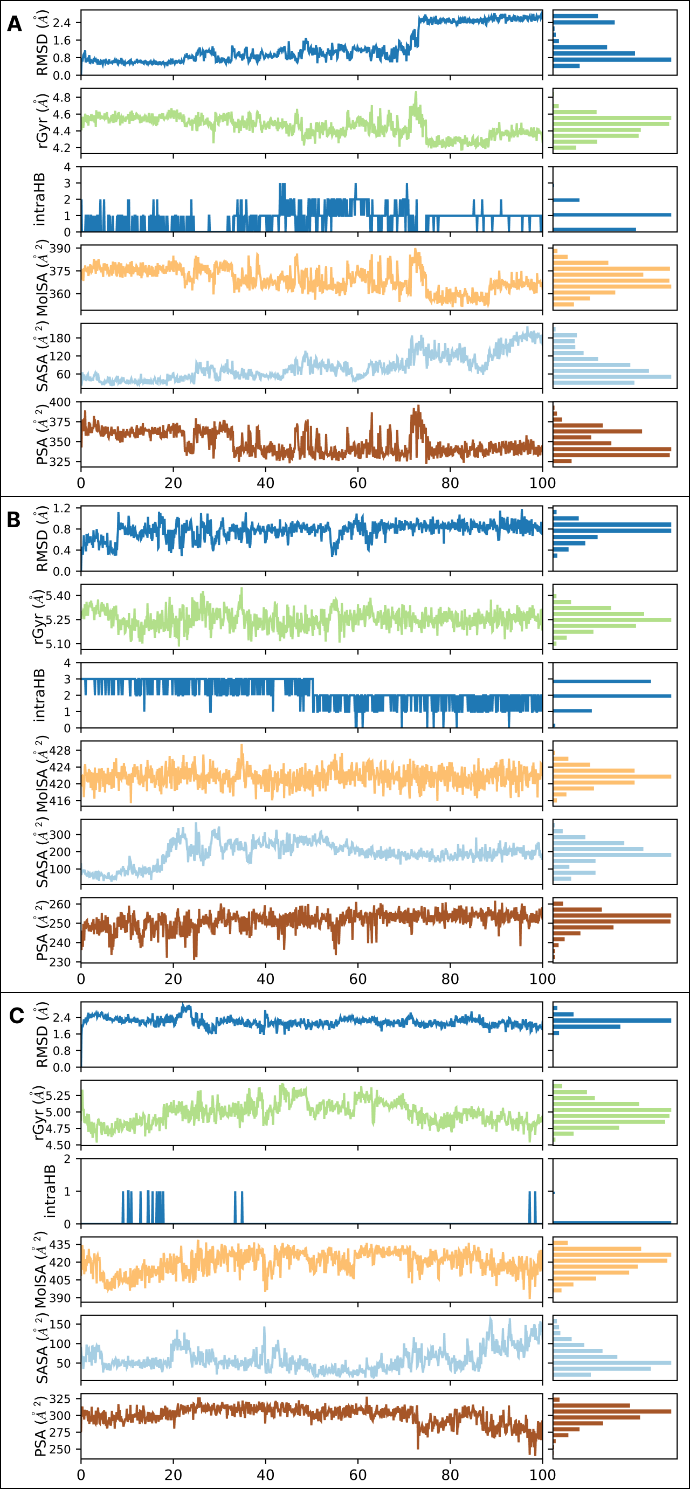

Figure 1 Variations in the properties RMSD, rGyr, MolSA, intraHB, SASA, and PSA for the three ligands (A:CNP0099279, B:CNP0298793, and C;CNP0347183) during 100ns of MD simulation time.
